# Complementation of a human disease phenotype in vitro by intercellular mRNA transfer

**DOI:** 10.1101/2024.11.06.622258

**Authors:** Gal Haimovich, Sandipan Dasgupta, Anand Govindan Ravi, Jeffrey E. Gerst

## Abstract

There is growing evidence that full-length mRNAs undergo intercellular transfer through long, thin cytoplasmic connections called tunneling nanotubes (TNTs), but whether transferred mRNAs are translated and effect cellular changes post-transfer is unknown. Using multiple lines of evidence, we show that transferred mRNAs undergo translation and can fully complement the phenotype of genetic mutations *in vitro*. For example, the human peroxisome biogenesis disorder, Zellweger Syndrome, results from mutations in genes like PEX5 and PEX6. We demonstrate that the co-culture of patient-derived PEX6 mutant fibroblasts or PEX5 knock-out cells with wild-type cells leads to *de novo* peroxisome biogenesis. Complementation occurs by TNT-mediated mRNA transfer and translation in acceptor cells and not by peroxisome transfer. We provide additional examples of genetic complementation via transfer of mRNAs encoding the HSF1 transcription factor or CRE recombinase. Our study provides evidence for the physiological significance of mRNA transfer and suggests a new approach for mRNA therapeutics.

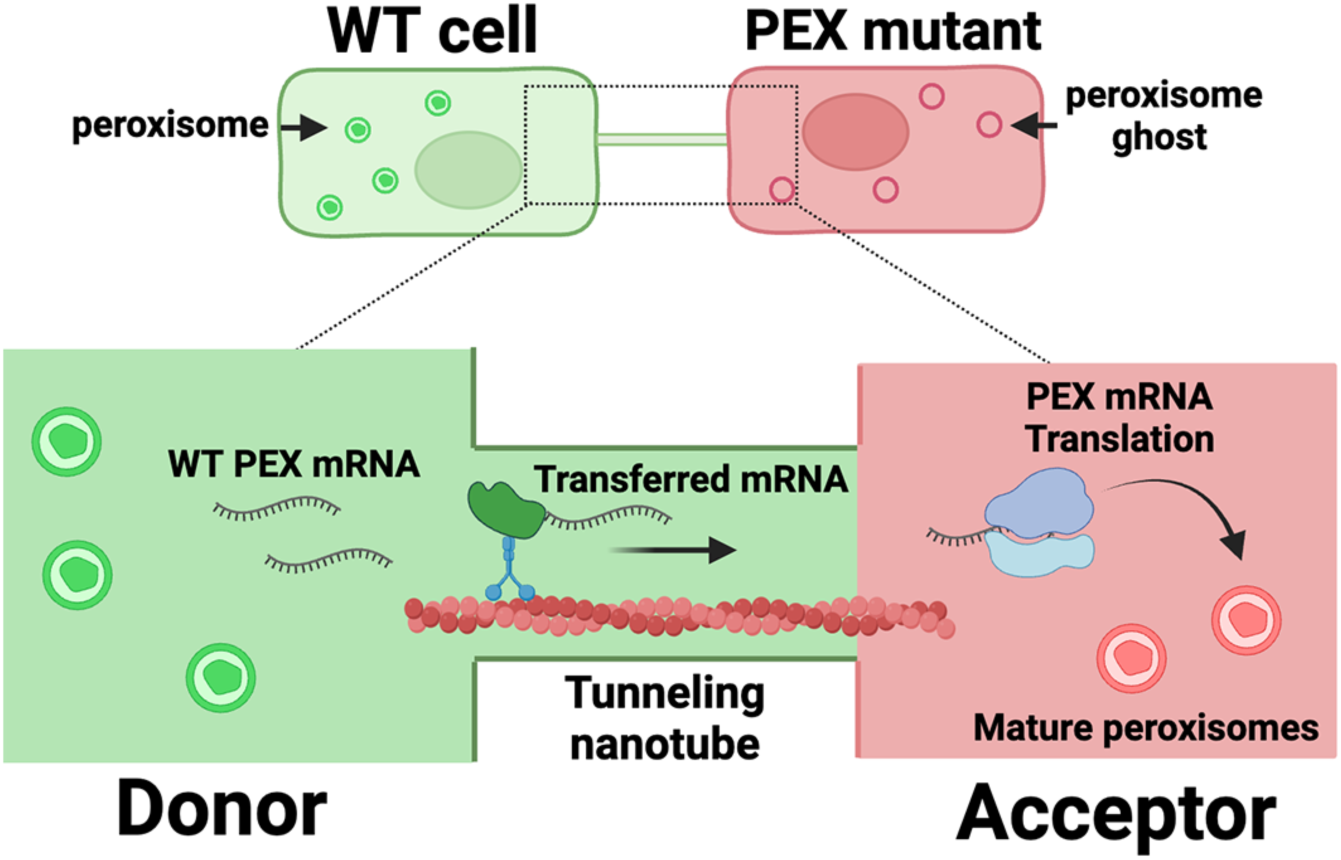

## Introduction

Messenger RNAs (mRNAs) have emerged as novel mediators of intercellular communication in a wide range of biological contexts, such as the propagation of viral infection, cancer cell metastasis, oxidative stress regulation, and immune cell maturation^1–6^. Once thought to undergo trafficking via secreted entities, such as extracellular vesicles or exosomes (referred collectively herein as EVs), few reports provide evidence that EV-transferred mRNAs are translated in and affect the physiology of recipient cells^7–9^. In contrast, we and others have shown that mRNAs are trafficked via direct intercellular connections, called tunneling nanotubes (TNTs; or membrane nanotubes)^3,10–16^. TNTs are long and thin cytoplasmic connections between cells, which are structurally distinct from lamellipodia and filopodia^17,18^. TNTs have been observed in many cell types and shown to transfer a wide range of cargoes, such as organelles, bacteria, viruses, proteins, and RNAs^3,19–23^.

We and others have shown by RNA sequencing (RNA-seq) that the range of transferred RNAs (the RNA transferome) correlates with gene expression levels in donor cells and constitutes ∼1% of each mRNA and lncRNA in the transcriptome^10,16,24^. Single molecule fluorescent *in situ* hybridization (smFISH) and live cell imaging studies have revealed that full-length mRNAs transfer via TNTs in a translation-independent, but expression level-, stress condition-and cell type-dependent manner^10,11,16^. In addition, transferred mRNAs are, perhaps, uniquely packaged^12^. As the transcriptome of recipient cells is strongly affected in co-culture experiments^10^, it suggested a possible contribution by the transferred RNAs to the cellular phenotype. Here, we examined whether transferred mRNAs are translated and induce phenotypic changes in recipient cells. We now show that the co-culture of cells lacking a functional mRNA with cells expressing the wild-type version can complement mutant phenotypes that result from the lack of the functional gene. For example, Heat-Shock Factor 1 (HSF1) is a transcription factor activated upon heat-shock (HS)^25^ and is required to induce the heat-shock response (*e.g.* HS-induced transcription of the heat-shock protein 70 (HSP70) gene, HSPA1A). While HSF1 knock-out (KO) cells are unable to properly induce the HS response, HSF1 mRNA transfer from donor cells can complement the deficiency, leading to re-activation of the HS response.

We also used an *in vitro* human disease model in which we co-cultured wild-type cells with SV40-transformed fibroblasts derived from a Zellweger Syndrome (ZS) patient that harbor a nonsense mutation in the PEX6 gene (referred to as PEX6mut cells hereafter)^26^. PEX6 encodes a predominantly cytosolic protein that belongs to the AAA family of ATPases that acts together with PEX1 as an export module of the PEX5 peroxisome protein import receptor^27^. Cells bearing mutations in the PEX5 or PEX6 genes have been reported to harbor “ghost” peroxisomes^28,29^ that are import-incompetent for some peroxisomal matrix proteins^29–31^. Importantly, we found that co-culture led to the *de novo* biogenesis of functional peroxisomes in the PEX6 mutant cells. Moreover, we observed the transfer of PEX6 mRNA from the donor cells to the mutants in a cell-to-cell, contact-dependent, manner and provide direct evidence for the *in situ* translation of transferred PEX6 mRNA.

Finally, we show that the transfer of CRE recombinase mRNA by either HEK293T cells or various mouse cell lines allows for DNA recombination in acceptor cells and the expression of a fluorescent protein reporter. Based on these findings, we conclude that TNT-mediated mRNA transfer can have a significant physiological impact on recipient cells and may be explored as a novel approach to gene therapy.

## Results

### HSF1 RNA transfer to HSF1^-/-^ cells and heat shock complementation

We first tested for the ability of mRNA transfer to complement the lack of HSF1. We obtained MEFs that originated from a HSF1 KO (HSF1^-/-^) mouse^32^. These cells do not induce the transcription of HSP70 upon HS, as compared to HSF1^+/+^ cells, but HS-induction of HSP70 in HSF1^-/-^ MEFs is restored upon overexpression of human HSF1 fused to GFP (HSF1-GFP) (**Supplementary Figure 1A-C**).

Next, we co-cultured HSF1^-/-^ MEFs with HSF1^-/-^ MEFs overexpressing HSF1-GFP for 48 hours. We found that HSF1-GFP mRNA transfers to the HSF1^-/-^ cells (**Supplementary Figure 1D**), as detected by smFISH and restores partial, albeit limited, heat shock responsiveness as exemplified by increased HSP70 transcription at transcription sites, without significantly increasing the number of transcription sites per cell (**Supplementary Figure 1E-G**). Although the results suggest that mRNA transfer can complement a deficiency, this system showed only a mild improvement over the KO, making it difficult to detect a physiological effect such as cell survival after heat-shock.

### Complementation of the peroxisome deficiency in Zellweger Syndrome cells by co-culture with wild-type cells

To test our hypothesis using an alternative system, we asked whether a co-culture of a peroxisome mutant cell line and a wild-type cell line can complement the mutant (peroxisome-deficient) phenotype. We designed a green fluorescent protein (GFP) reporter with a C-terminal peroxisome targeting signal type 1 (PTS1) consisting of serine-lysine-leucine, herein referred to as GFP-SKL. Expression of GFP-SKL led to a punctate pattern in wild-type HEK293T cells, which co-localized with the peroxisomal matrix protein, Catalase, and the PEX14 peroxisomal membrane protein, indicative of fully functional peroxisomes (**Supplementary Figure 2A and D**). We then tested SV40-transformed skin fibroblast cells derived from a ZS patient that harbor a nonsense mutation in PEX6 gene (c.1314-1321del (p.Glu439fs)) (see Methods section). This PEX6 mutation was previously described in other patients^33^. These cells are devoid of functional peroxisomes, as indicated by the diffuse staining pattern of Catalase (**Supplementary Figure 2B**), but still contain peroxisome ‘ghosts’, as indicated by the punctate pattern of PEX14 labeling (**Supplementary Figure 2E**). As a control, we stained HeLa cells that are CRISPR-generated KOs for PEX19 (ΔPEX19/PEX19^-/-^) cells. As expected^34^, these cells are also devoid of ‘ghosts’ as indicated by the diffuse staining of Catalase and the mitochondrial localization of PEX14 protein (**Supplementary Figure 2C and F**).

We generated a stable cell line of the PEX6mut cells expressing red fluorescent protein (RFP)-SKL and co-cultured it with HEK293T cells stably expressing GFP-SKL. There are several possible scenarios by which the complementation of peroxisomal biogenesis might possibly occur (**Figure 1A**). First, co-culture with the HEK293T-GFP-SKL cells could rescue the mutant peroxisome-deficient phenotype of PEX6mut cells by the transfer of PEX6 mRNA (or protein) and subsequent peroxisome biogenesis *de novo*. In this case, we expect to see red-colored peroxisomes in the acceptor cells due to the import of the cytoplasmic RFP-SKL into the restored peroxisomes. Second, we might observe green-labeled peroxisomes in these cells, which would indicate either cell-cell fusion or the transfer of whole GFP-labeled peroxisomes from the WT donors. Third, yellow peroxisomes could arise via the transfer of both PEX6 and GFP-SKL mRNA (or protein), or the transfer of whole peroxisomes from the donor cells, followed by the subsequent import of RFP-SKL protein into the green peroxisomes.

**Figure 1.**
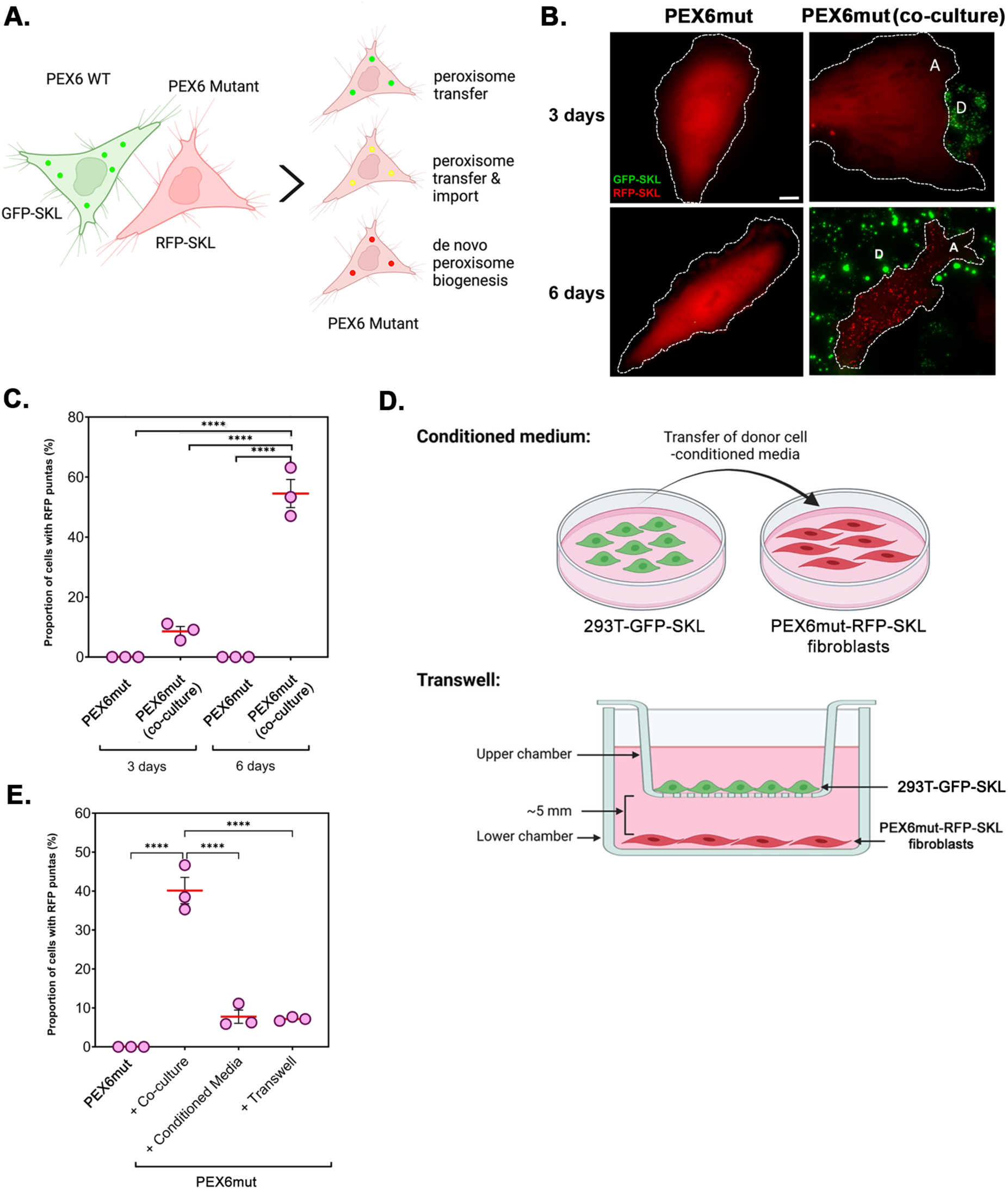
Complementation of PEX6 mutant cells in co-culture with WT cells. **A.** A pictorial depiction of the co-culture experiment: Donor HEK293T cells expressing a peroxisomal marker, GFP-SKL (containing green-colored peroxisomes) were co-cultured with acceptor PEX6mut cells expressing RFP-SKL for 3 days or 6 days. Complementation of peroxisomes could occur by *de novo* biogenesis in acceptor cells (leading to the formation of red-colored peroxisomes), whole organelle transfer (indicated by green-colored peroxisomes), or both (yellow-, green-and red-colored peroxisomes). **B.** Representative images of peroxisome complementation by live imaging after 3 or 6 days of mono-or co-culture. A: Acceptor cells, D: Donor cells. Dashed lines outline the approximate cell boundaries. Scale: 10µm. **C.** The proportion of cells with RFP-localized puncta in acceptor cells from three independent replicates is shown. Each circle represents one replicate (*e.g.* 30-35 cells per replicate). The horizontal red line and the error bars represent the mean and SEM, respectively. **D.** Schematic of the conditioned media and Transwell setup: Conditioned media was derived from a culture plate of 293T-GFP-SKL cells and transferred to a plate of PEX6mut-RFP-SKL cells (top illustration). In the Transwell setup, 293T-GFP-SKL cells were cultured in the top insert of a Transwell with 0.4µm perforations (bottom illustration). PEX6mut-RFP-SKL cells were cultured in the lower chamber of a 12-well glass bottom plate. Donor and acceptor cells were separated by ∼5mm. **E.** The proportion of cells with RFP-localized puncta in acceptor cells, as derived from three independent replicates is shown. Each circle represents one replicate (30-40 cells per replicate). The horizontal red line and error bars represent the mean and SEM, respectively.

Whereas RFP-SKL remained cytoplasmic in PEX6mut cells cultured alone (**Figure 1B**, left panels), we observed that red-labeled puncta appear in about half of the PEX6mut cells after six days of co-culture with donor cells (HEK293T GFP-SKL), but rarely earlier (**Figure 1B-C**). These puncta were similar in size to those observed in wild-type cells labeled for peroxisome localization (**Supplementary Figure 2A and D**). Importantly, we did not detect any green-or yellow-colored peroxisomes in the acceptor cells, indicating that the complementation is not mediated by whole organelle transfer, but likely *via* the transfer of PEX6 mRNA and/or protein, and occurs in a time-dependent manner (**Figure 1B**). These results indicate *de novo* peroxisome biogenesis.

Although PEX6 mRNA was not detected in isolated HEK293T-derived exosomes^35^, we tested if indeed the observed complementation was mediated by mechanisms that required cell-to-cell contact, such as TNTs, or by diffusion, such as EVs. We separated the donor and acceptor cells in space by culturing them in two distinct chambers of a Transwell setup or culturing the acceptor cells with conditioned culture medium collected from donor cells (**Figure 1D**). The number of acceptor cells with RFP-labeled peroxisomes was highest under co-culture conditions and was ∼10-fold less in the Transwells or when cultured with donor cell-conditioned media (**Figure 1E**). This clearly indicates that the mechanism involved in peroxisome complementation requires direct cell-to-cell contact.

### PEX6 mRNA transfers in co-culture

We next determined if PEX6 mRNA transfers from donor HEK293T cells to the PEX6mut cells. We could not use smFISH to detect the transfer of endogenous wild-type PEX6 mRNA from HEK293T cells since the transferred mRNA cannot be differentiated from the mutant endogenous PEX6 mRNA due to their close identity. Thus, we constructed a reporter construct whereby the gene for the Tomato red fluorescent protein was fused to the 3’end of the PEX6 gene, upstream of the native untranslated region (UTR), and expressed under a constitutive Ubiquitin C promoter (**Supplementary Figure 3A**). To check the ability of the construct to rescue the mutant phenotype of PEX6mut cells, we transiently transfected PEX6-Tomato together with GFP-SKL. In this case, we observed robust complementation of the PEX6mut phenotype within three days post-transfection, as evidenced by the punctate pattern of GFP-SKL (**Supplementary Figure 3B**). In contrast, we did not observe complementation when the mutant cells were transfected with a construct expressing tandem Tomato (tdTomato) gene together with GFP-SKL (**Supplementary Figure 3B**).

We then generated a HEK293T donor line that stably expresses PEX6-Tomato at levels approximately 25-fold more than native PEX6 mRNA, relative to parental HEK293T cells (**Supplementary Figure 3C**). To detect the transfer of PEX6-Tomato mRNA, we co-cultured PEX6mut cells with PEX6-Tomato expressing cells and visualized the mRNA by smFISH using probes specific for the Tomato sequence (**Figure 2A**). We observed that PEX6-Tomato HEK293T cells express on average 300 copies of the mRNA after 2 days in co-culture, while this decreases by half after 4 days. Importantly, we observed Pex6-Tomato mRNA transfer to PEX6mut cells, with approximately 40 copies per cell detected after 2 days of co-culture, increasing to 250 copies per cell within 4 days of co-culture (**Figure 2B**). As seen for peroxisome biogenesis complementation, PEX6-Tomato mRNA transfer is solely dependent on cell-cell contact (**Figure 2C** and **Supplementary Figure 4**).

**Figure 2.**
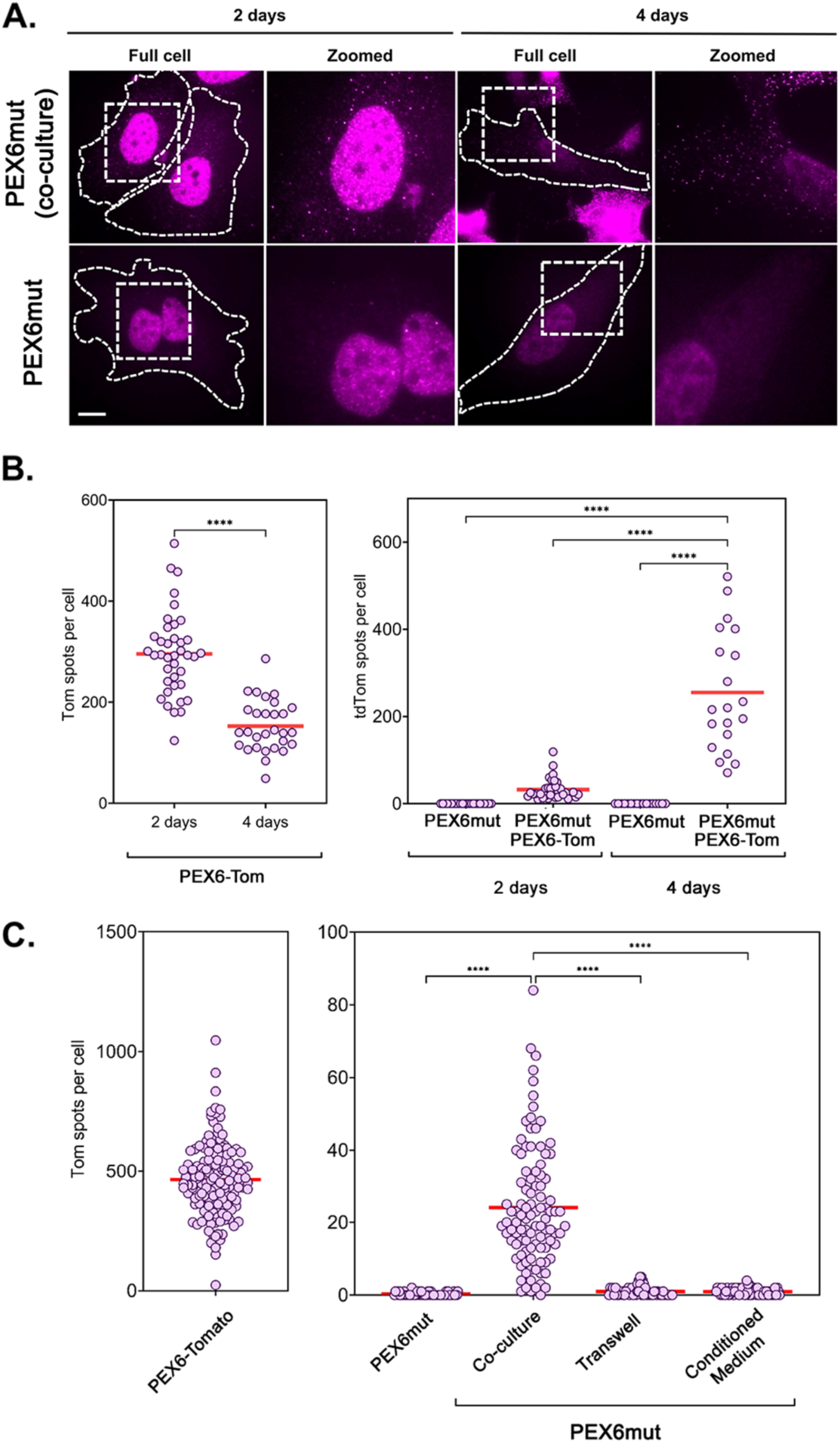
Intercellular transfer of PEX6 mRNA from PEX6-Tom donor cells to PEX6mut cells. **A.** Representative smFISH images depicting the presence of PEX6-Tomato mRNA in PEX6mut acceptor cells cultured alone (bottom panels) or in co-culture with HEK293T cells overexpressing PEX6-Tomato (top panels) for 2 days and 4 days, respectively. A single spot corresponds to a single molecule of PEX6-Tomato RNA, as detected by smFISH for tdTomato. Rectangles marked by dashed boundaries are zoomed. Scale: 10 µm. (**B-D**) Distribution of PEX6-Tomato mRNAs in donor and acceptor cells under different conditions. Each dot represents the number of mRNAs in a single cell. The red bar represents the mean of each distribution. ****: P<0.0001. **B.** Distribution of PEX6-Tomato mRNAs in acceptor cells in co-culture with donor HEK293T-PEX6-Tomato cells for either 2 or 4 days. **C**. Distribution of PEX6-Tomato mRNAs in either donor cells (left panel) or in acceptor cells (right panel) cultured alone, in co-culture with donor cells, with donor cell-conditioned media, or in close proximity to donor cells using Transwell inserts. See Figure 1D for a schematic of the conditioned media and Transwell setup. Representative smFISH images are shown in **Supplementary Figure 4**.

### Transferred PEX6 mRNA undergoes translation in acceptor cells

We next determined whether transferred PEX6-Tomato mRNA is translated in acceptor cells. We inserted a 7xHA tag at the 5’ end of the PEX6-Tomato gene and created stably expressing donor cells as before. We used two strategies to detect translation. First, we co-cultured the donor and acceptor cells for two days, then performed smFISH-immunofluorescence (FISH-IF) using fluorescent probes for Tomato and antibodies against the HA Tag (**Figure 3A**). Colocalized/adjacent signals of the smFISH and IF spots would indicate that the transferred mRNA is being translated, whereas separate signals would suggest that the protein was transferred independently of the mRNA (**Figure 3B**). We observed robust transfer of 7xHA-PEX6-Tomato mRNA from donor cells to PEX6mut acceptor cells (**Figure 3C**) and could detect HA spots in very close proximity to Tomato mRNA spots (**Figure 3D-E**). Importantly, the two signals were separated by ∼300-400 nm (**Figure 3E-F**), which is even shorter than the predicted linear distance between the two signals of PEX6-Tomato mRNA (*i.e.* ∼1000nm, assuming 0.33 nm/nucleotide^36^ of a ∼3000nt mRNA). Overall, we observed that ∼40 molecules of mRNA per cell transferred to the acceptor cells, out of which around 10% were found adjacent to HA spots, thus, being likely candidates for translation sites (**Figure 3C and G**). An alternative, though unlikely explanation is that PEX6 protein from donor cells undergoes transfer together with its mRNA. Quantification of the average total cell IF signal intensity revealed that the amount of total HA-tagged protein in acceptor cells in co-culture is significantly higher than that of acceptor cells alone (*i.e.* background) (**Figure 3H**). This can arise either due to translation of the transferred mRNA or transfer of sufficient quantities of HA-tagged protein.

**Figure 3.**
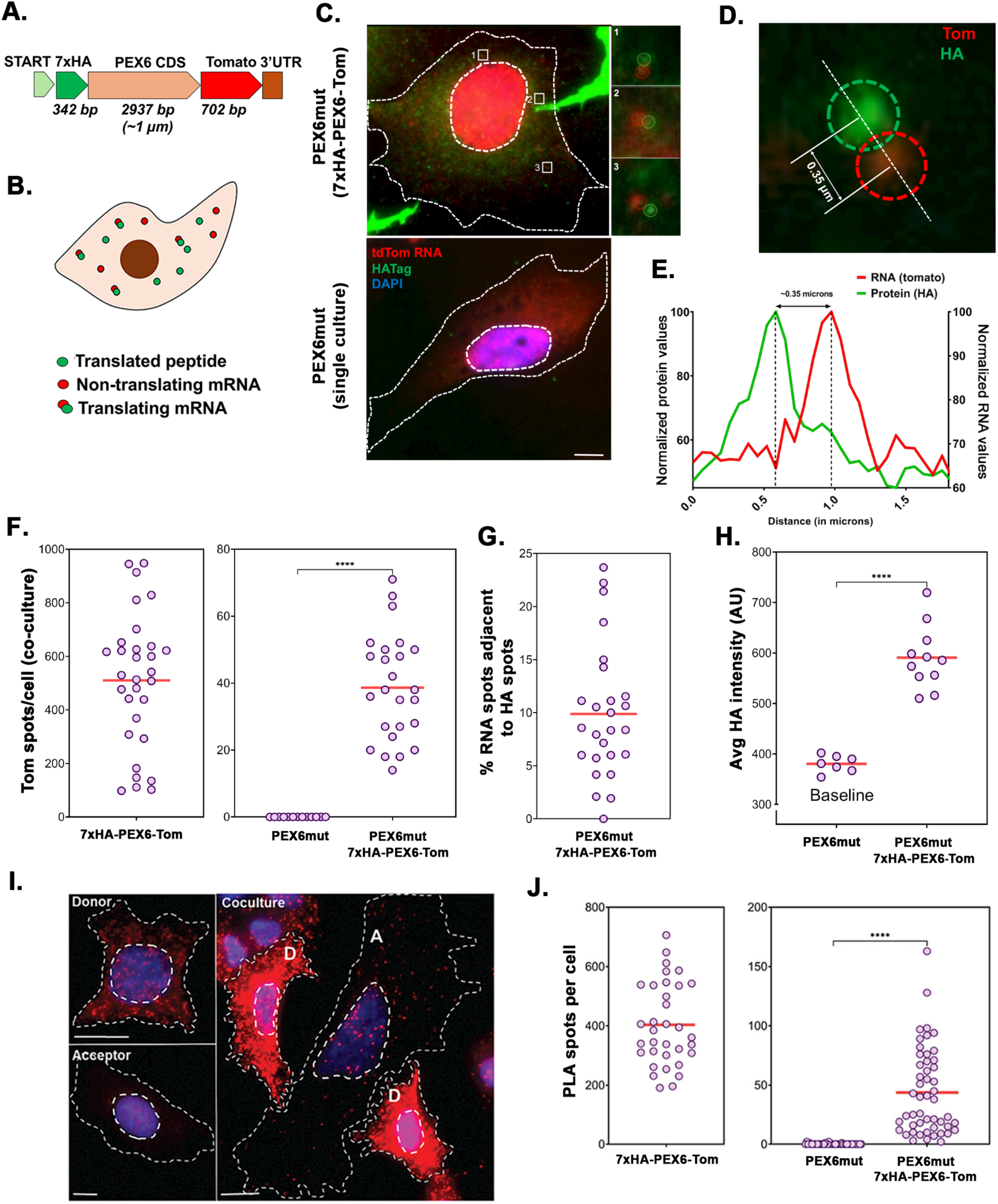
Translation of transferred PEX6 mRNA. **A.** Schematic design of the PEX6 construct used to detect translation sites: a 7xHA tag was inserted in-frame to the 5’ end of the PEX6 CDS; the gene for Tomato was inserted in-frame at the 3’ end of PEX6, which abolishes the PEX6 termination signal, and upstream of the PEX6 3’UTR. Lengths of each segment are indicated. Based on structural studies, the linearized mRNA of PEX6 CDS in likely to be ∼1 µm in length. START: Start codon; STOP: Stop codon. **B.** Schematic of expected outcomes of FISH-IF: If transferred mRNAs undergo translation, then FISH spots from Tomato mRNA (red) and IF spots from 7xHA Tag (green spots) are expected co-localize or be in close proximity to each other. Single green spots correspond to the mature protein and single red spots correspond to non-or post-translated mRNAs. **C**. Representative smFISH-IF Images. PEX6mut (acceptor) cells were cultured either alone (top) or with donor HEK293T cells expressing 7xHA-PEX6-Tomato for 2 days (bottom). Cells were fixed and smFISH (red color) and IF (green color) were performed sequentially. Three sites with adjacent FISH and HA spots are outlined with white squares and zoomed in. Dashed lines outline the approximate cell and nuclear boundaries. Scale: 10 µm. **D.** An example of adjacent FISH-IF spots is shown. The Tomato FISH spot is shown in red and HA IF spot in green. Dashed green and red circles outline the approximate perimeters of the IF and FISH spots, respectively. The distance between the centers of the two spots is indicated, as measured by ImageJ. **E**. Intensity quantification of the image shown in D. The intensity profiles of the IF and FISH signals were measured and plotted along the dashed line which transverses the spots using the *plot profile* function of ImageJ. The values were then normalized so that the peaks are at 100 a.u. for each channel and the two plots combined on the same graph. **F.** Distribution of HA-PEX6-Tomato mRNAs in donor and acceptor cells: Donor (293T-HA-PEX6-Tomato) and acceptor cells (PEX6mut) were either cultured alone or together in co-culture for 2 days and HA-PEX6-Tomato mRNA was detected using FISH for Tomato mRNA in donor cells (left panel), acceptor cells alone or acceptor cells in co-culture (right panel). Each dot represents the number of mRNAs in a single cell. The red bar represents the mean of each distribution. ****: P<0.0001. **G**. Percentage of transferred PEX6-Tom RNAs in co-cultured acceptor cells found in close proximity with an IF spot (<0.5 µm). Each spot represents measurements from a single cell. Red bar shows the mean of the distribution. **H**. Intensity quantification of HA expression: The HA Tag was detected by IF in either acceptor cells alone (PEX6mut) or in acceptor cells co-cultured with PEX6-Tom donors and the measured fluorescence intensity plotted. Red bar shows the mean of the distributions. ****: P<0.0001. A.U. -arbitrary units. **I.** Maximum projection images of puro-PLA of PEX6mut (acceptor) cells that were cultured alone or with donor HEK293T cells expressing 7x-HA-PEX6-Tomato for 2 days. Red spots indicate PLA products. Blue: DAPI. D: Donor; A: Acceptor; Scale: 10 µm. **J.** The graph shows the distribution of the number of PLA spots in 7xHA-PEX6-Tom donor cells (left panel) and PEX6mut acceptor cells cultured either alone or in co-culture with 7xHA-PEX6-Tom donors (right panel). Each spot represents a single cell and the red indicates the average number of spots. ****: P<0.0001

To verify the possibility of post-transfer mRNA translation, we performed Puromycin Proximity Ligation Assay (Puro-PLA)^37^ to detect translation *in situ*. Briefly, following the co-culture of acceptor PEX6mut cells with donor 7xHA-Pex6-Tomato cells, the cells were treated with puromycin which causes premature translation termination and releases nascent puromycinylated polypeptides tagged at the C-terminus. Thus, *in situ* translation of PEX6 mRNA can be detected using anti-HA and anti-puromycin antibodies followed by PLA, where each nascent peptide is detected as a spot (**Figure 3I**). The presence of these spots in acceptor cells in co-culture is direct evidence of nascent translation of the transferred PEX6-Tomato mRNAs, since transferred mature proteins are not puromycinylated and thus not detected. Indeed, an average of 45 PLA spots per cell was observed in PEX6mut cells after 2 days of co-culture with donor HA-Pex6-Tomato cells, while no spots were detected in the acceptor cells alone (**Figure 3J-K**). We also detected nuclear-associated PLA spots, which could originate in maximum projection images from the cytoplasm either above or below the nucleus (**Supplementary Figure 5**) or perhaps from the diffusion of small puromycinylated peptides to the nucleus prior to or during fixation, or even from non-specific binding^38^.

### Intercellular mRNA transfer can complement other peroxisome biogenesis defects

To test whether we can restore peroxisomes in another peroxisome biogenesis mutant, we obtained a PEX5 knock-out cell-line derived from HEK293T-rex cells (*i.e.* PEX5^-/-^). We co-cultured RFP-SKL expressing PEX5^-/-^ cells with GFP-SKL expressing WT cells and counted the number of cells exhibiting RFP foci after six days. Co-culture with WT cells led to a ∼2.5-fold increase in the number of peroxisome-positive cells, as compared to PEX5^-/-^ cells cultured alone (**Supplementary Figure 6**).

### Activation of lox-STOP-lox-tdTomato gene expression by CRE mRNA transfer *in vitro*

To study this process further, we developed a third model that employs the Ai9 mouse line^39^ which bears a LoxP-flanked STOP cassette (*i.e.* having stop codons in all three reading frames and a triple poly-A signal) located upstream of a genome-integrated tdTomato open reading frame (ORF). We obtained MEFs from this mouse and immortalized them by expression of SV40 large T antigen. Next, we created lentiviral vectors expressing CRE recombinase from the EF1α promoter; one vector that co-expresses GFP and another that co-expresses a tandem MS2 coat protein fused to GFP (tdMCP-GFP). In the 3’UTR of CRE we inserted multiple MS2 stem loops to allow detection of the mRNA by FISH or to inhibit mRNA transfer by co-expressing tdMCP-GFP^11,40^. Expression of CRE recombinase in the Ai9 MEFs removes the STOP cassette, allowing the cells to express tdTomato and the cells turn red (**Supplementary Figure 7A**).

We transduced HEK293T cells with our two CRE constructs or GFP alone. CRE protein and mRNA levels in the donor cells expressing either CRE/GFP or CRE/tdMCP-GFP were similar, indicating tdMCP-GFP did not affect mRNA or protein expression therein (**Supplementary Figure 7B-C**). We then co-cultured the Ai9 MEFs with HEK293T cells that expressed CRE/GFP, CRE/tdMCP-GFP or GFP alone. After 72 hrs, we found that ∼1-3% of the MEFs were tdTomato positive as compared to 0% in the GFP control (**Figure 4A**). This result indicates that CRE recombinase mRNA underwent transfer, translation, and removed the STOP cassette in these cells by recombination. Interestingly, less tdTomato positive cells were seen when CRE was co-expressed with tdMCP-GFP in the donor cells, which strengthens our hypothesis that CRE mRNA, and not Cre protein, is transferred (**Figure 4A**). Indeed, smFISH analysis confirmed that less CRE-MS2 mRNA was transferred when co-expressed with tdMCP-GFP (**Figure 4B**). Similar results were obtained with wild-type MEFs expressing CRE/GFP or CRE/tdMCP-GFP as donors (**Figure 4C**).

**Figure 4.**
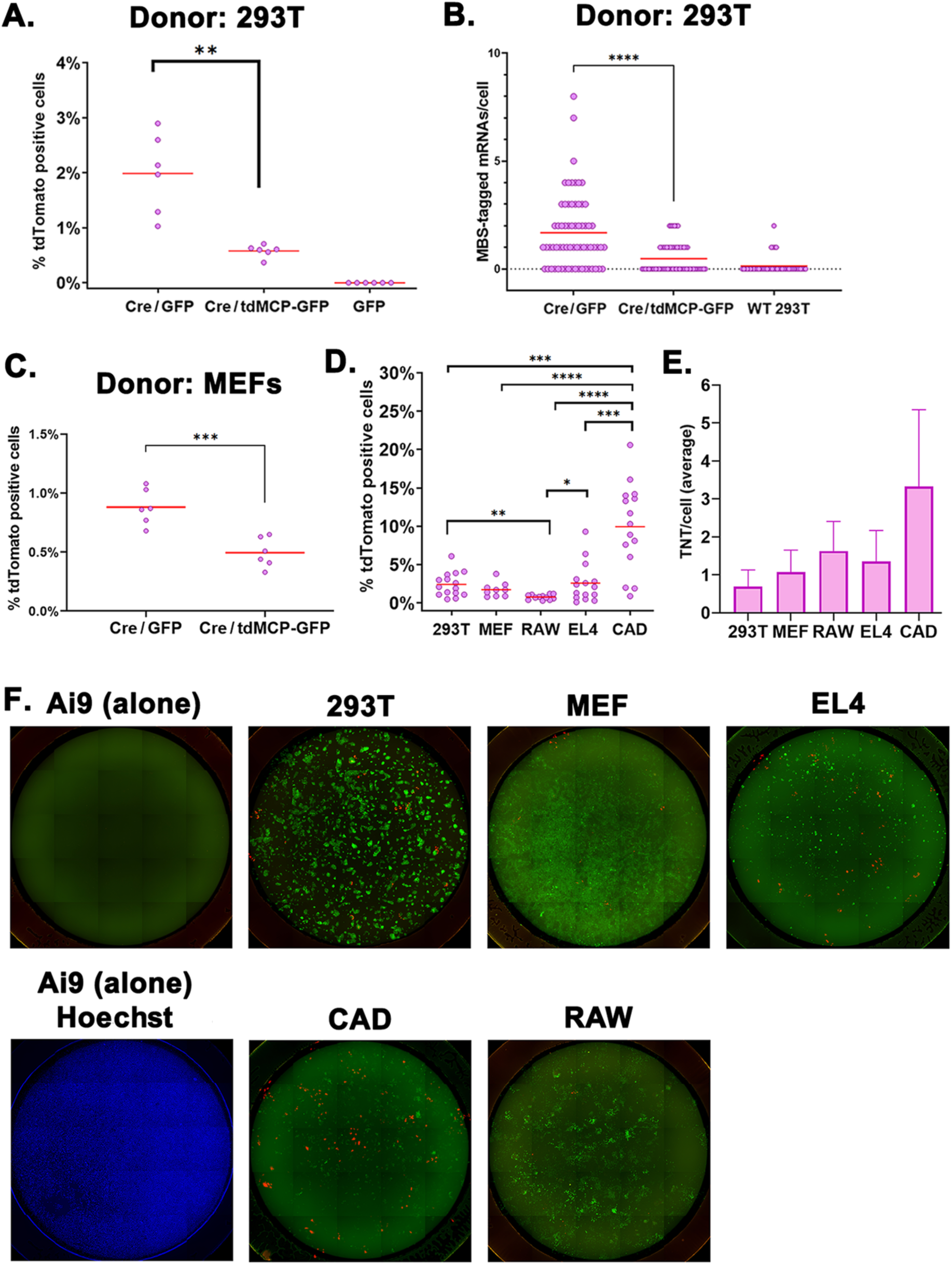
Activation of tdTomato expression by CRE mRNA transfer. **A.** Percentage of Ai9 tdTomato-positive cells after 3 days of co-culture with CRE-expressing HEK293T cells. Each dot represents one biological repeat of the experiment. **B.** CRE mRNA transfer to Ai9 cells from HEK293T cells expressing CRE/GFP, CRE/tdMCP-GFP, or control cells. Each dot represents a single cell. **C.** Percentage of Ai9 tdTomato-positive cells after 3 days of co-culture with CRE-expressing MEFs. Each dot represents one biological repeat. **D.** Percentage of Ai9 tdTomato-positive cells after 3 days of co-culture with various cell lines expressing CRE. Each dot represents one biological repeat. Note that panel D includes data from panels A and C. **E.** The average number of TNTs per cell connecting donor to acceptor cells after 24hr in co-culture. Note that only donor-acceptor, but not donor-donor or acceptor-acceptor TNTs were counted. For all panels**: * -** p<0.05; ** -p<0.01; *** p<0.001; **** - p < 0.0001. **F. Images of donor cells co-cultured with Ai9 cells.** Compound images of the 25 fields that comprise one entire well of a 96-well plate are shown. Merge of the green (GFP) and red (tdTomato) channels is shown; for Ai9 cells alone the Hoechst (blue) channel is shown as proof of the presence of cells in the well. Donor cell lines expressing CRE/GFP are noted above the images. Note that the GFP expression levels differ between donors (similar to the variations seen with CRE; (**Supplementary Figure 7D**).

We then compared the efficiency of various cell types to confer mRNA transfer using the Ai9 cell tdTomato activation assay: HEK293T, MEFs, EL4 (T lymphoblasts), RAW267.4 (macrophages) and CAD (neuron-like) cells. All donor cell types tested could turn Ai9 cells red. HEK293T, MEFs and EL4 cells showed similar efficiency, whereas RAW267.4 cells were less efficient and CAD cells were the most efficient (**Figure 4D and F**). Consistent with previous results, there were no red cells observed (0%) when the Ai9 MEFs were co-cultured with any of the CRE donor cells using the Transwell set up. Thus, the STOP segment is removed by recombination only when donor and recipient cells are in direct contact and when the mRNA can freely transfer through TNTs. mRNA transfer efficiency appeared to be affected by both mRNA expression levels and the level of TNT formation. We found a wide variability of the mRNA levels amongst the donor cells, despite the fact the same promoter was used (**Supplementary Figure 7D**). This correlated with the number of transferred mRNAs detected in co-cultures (**Supplementary Figure 7E**), but not with the percentage of tdTomato-positive cells (**Figure 4D**). We also counted the number of connecting TNTs and found that most donors produced similar levels of donor-acceptor TNTs, except CAD cells, which produced 2-3-fold more (**Figure 4E and Supplementary Figure 8**). This increase in TNT levels correlated with their higher efficiency in activating tdTomato expression in Ai9 cells.

## Discussion

In this study, we used imaging-based approaches to determine if intercellular mRNA transfer from wild-type cells can complement a deleterious mutation in mutant cells. We found that the co-culture of wild-type cells with a peroxisome-deficient ZS patient-derived cell line mutated for the PEX6 gene rescues the mutant phenotype (**Figure 1**). We also found that PEX6-Tomato fusion mRNA transfers from an overexpressing donor cell line to PEX6mut acceptor cells via direct cell-to-cell contact (**Figure 2**). The level of mRNA transfer is higher than we reported previously for other mRNAs^10,11^, which is likely explained by the prolonged co-culture conditions employed. Importantly, we demonstrate that transferred mRNAs are translated in acceptor cells (**Figure 3**). A recent pre-print showed indirect evidence for translation of transferred mRNAs^24^. However, to the best of our knowledge, ours is the first study to directly demonstrate translation *in situ* of a transferred mRNA into protein in acceptor cells. This is in addition to indirect evidence showing that the transferred mRNAs are translated into functional proteins by observing changes in cell physiology (**Figures 1** and **4** and **Supplementary Figures 1 and 6**).

Complementation by intercellular mRNA transfer is not unique to PEX6 mRNA. We showed that this mechanism complements lack of the PEX5 peroxisomal import receptor in PEX5^-/-^ cells (**Supplementary Figure 6**) and lack of the HSF1 transcription factor in HSF1^-/-^ cells (**Supplementary Figure 1**). Furthermore, it can induce genomic recombination events in cells via the transfer of CRE recombinase mRNA to Ai9 cells (**Figure 4** and **Supplementary Figure 7**). However, these systems appeared to be less efficient than PEX6 complementation for reasons yet to be determined. Gene complementation by intercellular RNA transfer appears slow and time-dependent over the course of days. This implies that it requires sufficient time to build up a critical mass of protein from transferred mRNAs to affect phenotype of acceptor cells. Interestingly, we did not find transfer of intact peroxisomes although nanotubes were shown to transport other organelles, such as mitochondria, lysosome, endoplasmic reticulum and vesicles^17,19,41–44^. Although some proteins were shown to transfer via TNTs^41,45,46^, our results with MS2-tagged CRE mRNA in MEFs and HEK293T cells strongly suggest that mRNA (and not protein) transfer confers phenotypic changes to acceptor cells. Our results using CAD cells, which yielded the highest level of tdTomato gene activation (**Figure 4D**), but nearly undetectable levels of transferred mRNA (**Supplementary Figure 7E**), provides a conundrum yet to be resolved. This might result from both protein and mRNA transfer or from the high translatability of the few transferred mRNAs. However, it is clear that the phenotypic changes observed were due to contact-dependent processes (**Figure 1E**). Notably, we did not observe any red Ai9 cells when CRE mRNA donor cells were separated from the tdTomato acceptors using Transwells. Moreover, tdTomato activation correlated directly to the number of TNTs per cell (**Figure 4E**).

Our results provide the first strong evidence that transferred RNAs are functional and undergo translation in the acceptor cells (**Figure 3**). We validated the finding by directly visualizing the nascent translated proteins derived from transferred PEX6 mRNAs by a proximity ligation approach^37^. Interestingly, smFISH results obtained using different fixation methods previously suggested to us that transferred mRNAs might be packaged with an unknown protein coat or capsule^12^. If so, this could necessitate removal to allow for translation of the transferred mRNA and perhaps explain why our FISH-IF results suggest that only ∼10% of the transferred mRNAs are translated at any given time (**Figure 3G**). Overall, our results suggest a model by which an mRNA can transfer from donor cells to acceptor cells in co-culture, undergo translation, and complement a genetic deficiency (**Figure 5**).

**Figure 5.**
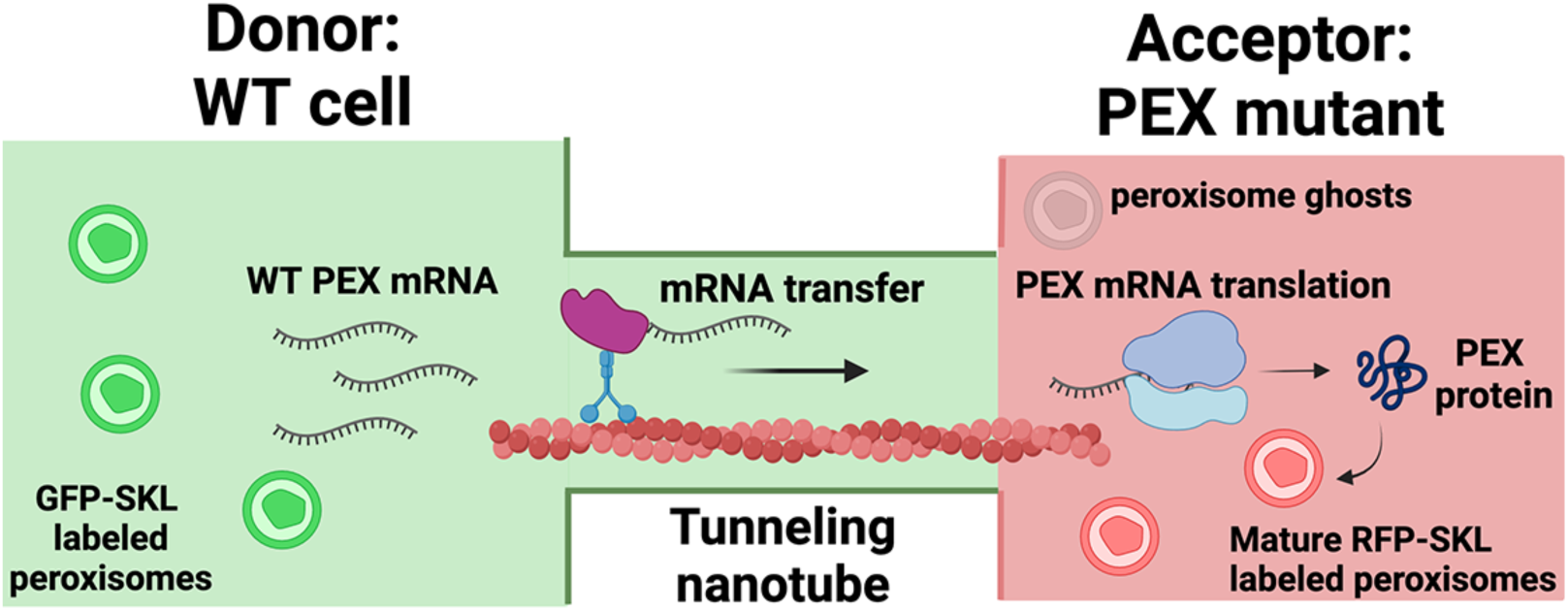
Schematic model for the reconstitution of peroxisome biogenesis in PEX mutant cells upon mRNA transfer. Our model for intercellular mRNA exchange (ImAx) suggests that wild-type (WT) mRNAs transfer from WT cells to mutant cells via tunneling nanotubes (TNTs), probably via a molecular motor (*e.g.* myosin) along actin filaments^11^. Once delivered to the acceptor cells, the transferred mRNA undergoes translation and the expressed protein is able to directly affect cell physiology. In the example shown, PEX mutant acceptor cells expressing RFP fused to a peroxisome targeting signal (RFP-SKL) show only incomplete/non-functional peroxisomes (*e.g.* unlabeled peroxisome ghosts). Upon co-culture of the acceptor cells with wild-type (WT) cells both expressing GFP-SKL (green-colored peroxisomes) and the appropriate PEX gene, we observed *de novo* peroxisome biogenesis to form mature RFP-labeled peroxisomes. Peroxisome biogenesis in the acceptor cells correlated with the transfer of PEX mRNA and PEX protein translation. Peroxisomes were not observed to undergo intercellular transfer, while RNA transfer was found to be cell contact-dependent and exosome-independent.

It should also be noted that the PEX6mut and PEX5 KO acceptor cells harbored peroxisomal “ghosts”^28,29^ (**Supplementary Figure 2**), which are incompetent for import of some, but not all peroxisomal matrix proteins^28,29^. Complementation of these mutants may be easier than for cells lacking PEX genes required for early peroxisomal biogenesis, such as PEX19 or PEX3, which are involved in peroxisome membrane formation^47^. Indeed, earlier attempts to complement a PEX19 KO mutant in our lab failed to show peroxisomal biogenesis after a week of co-culture. Previous complementation of peroxisome mutations have been primarily achieved by transfection of the cDNA of PEX genes into cells or by cell fusion with healthy fibroblasts^48^. In contrast, our study provides a novel approach of complementing peroxisome mutant phenotype by transfer of functional PEX mRNAs from a neighboring cell.

The intercellular trafficking of organelles, mainly mitochondria and lysosomes, from pluripotent stem cells, has been gaining traction as a novel modality for correcting genetic defects where mitochondrial or lysosomal function was found to be lacking^19,49,50^. A recent pre-clinical study showed that the injection of mesenchymal stem cells into the eyes of *Ndusf4-*KO mice could rescue mitochondrial function and corresponding degeneration of retinal ganglion cells^49^. However, our results show that whole peroxisomes do not transfer via TNTs or other mechanisms. Thus, Zellweger Syndrome defects must be rectified by other mechanisms. There are at least two pre-clinical reports of rAAV mediated delivery of the PEX1 gene to the eyes of a PEX1-G844D mouse model to rescue the retinopathy of the affected mice^51,52^. Our data suggest that the administration of PEX gene over-expressing cells (*e.g.* hematopoietic stem cells) could be a strong candidate approach to rescue ZS patients.

Interestingly, allogeneic hematopoietic stem cell transplantation (HSCT) was successful in mitigating the main phenotype of a mild ZS patient^53^. In addition, genetically modified HSCT for the treatment of X-linked adrenoleukodystrophy (X-ALD; a peroxisomal disorder) and metachromatic leukodystrophy (MLD; a lysosomal storage disease) were recently FDA-approved (Skysona^TM^) and EMA/FDA-approved (Libmeldy^TM^), respectively. Consistent with our model of complementation via TNT-mediated mRNA transfer, TNT formation was required for successful bone marrow transplantation treatment of a mouse model for Dent disease (a rare X-linked kidney tubulopathy)^54^. In addition, HSCT that over-expressed the PPT1 gene in a mouse model for infantile neuronal ceroid lipofuscinosis, a lysosomal storage disorder, improved the physiological condition of the mice better than HSCT using endogenously expressed PPT1^55^. This is consistent with our finding that higher gene expression levels in donor cells leads to more mRNA transfer^10,16^.

Work from various labs over the past 20 years has shown that many types of immune cells form tunneling nanotubes (TNTs), which form direct connections between cells^56–62^. Indeed, we show that immune cell lines, such as EL-4 and RAW264.7, form TNTs with acceptor MEFs and can transfer the CRE mRNA to elicit a phenotypic change in Ai9 cells. Moreover, a recent preprint suggested that macrophages might transfer endogenous mRNAs *in vivo*, based upon the analysis of previously published single-cell RNA-seq data of human xenografts in mice ^63^. While the *in vivo* work was not validated and no mechanism suggested, the authors do show that RAW264.7 can transfer FTL1 mRNA in an *in vitro* co-culture setup to HEK293T and HeLa cells, which confirms our current findings that macrophages can transfer mRNAs.

Thus, we predict that HSCT is successful due to the ability of HSC’s differentiated progeny (*i.e.* innate and adaptive immune cells) to transfer the wild-type mRNA to all affected cells and tissues throughout the body and thus alleviate the disease phenotype. Accordingly, HSCT may prove highly useful in the treatment of recessive human genetic disorders and our simple *in vitro* model of CRE-mediated activation of Ai9 STOP-tdTomato cells by mRNA transfer can be leveraged to improve transfer and thereby enhance the means of treatment.

## Materials and Methods

### Cell lines and Culture Conditions

HEK293T cells were purchased from ATCC. Immortalized wild-type (WT) MEFs were previously described^11^. Immortalized patient-derived PEX6mut^26^ and CRISPR knock-out HEK T-REx293 PEX5 cells^29^ were a kind gift from Ralf Erdmann (Ruhr Universitat Bochum, Germany). CRISPR knock-out ΔPEX19 HeLa cells ^34^ were a kind gift from Ron Kopito (Stanford University, CA USA). Immortalized HSF^-/-^ MEFs and isogenic WT MEFs (HSF1^+/+^)^32^ were a gift from Ruth Scherz-Shouval (Weizmann Institute of Science, Israel).

Ai9 MEFs were extracted from E13.5 embryos of Ai9 strain (B6;129S6-Gt (ROSA)26Sor^tm9(CAG-tdTomato)Hze/^J (Jackson Strain #007905) (a gift of Steffen Jung, Weizmann Institute of Science, Israel) and immortalized by transfecting the cells with plasmid pBSSVD2005 expressing SV40 large T antigen (a gift from David Ron (University of Cambridge, UK); Addgene plasmid #21826) according to a previously published protocol^64^. Cells were passed continuously and the colonies that survived after subsequent passages were considered immortalized and mixed together.

PEX6mut fibroblasts, PEX19^-/-^ KO HeLa cells, PEX5^-/-^ KO cells, HEK293T and MEF cell lines were cultured in DMEM (high glucose), supplemented with 10% Fetal Bovine Serum (FBS), 1 mM sodium pyruvate and antibiotics (0.1 mg/mL streptomycin and 10 U/mL penicillin). CAD cells^65^ were a gift from Chiara Zurzolo (Institute Pasteur, France). The cells were cultured in DMEM/F-12 1:1 media supplemented with 10% Fetal Bovine Serum (FBS), 2mM L-glutamine, 1 mM sodium pyruvate and antibiotics. EL4 cells (ATCC TIB-39) were a gift from Benjamin Geiger (Weizmann Institute of Science, Israel). The cells were cultured in DMEM (high glucose), supplemented with 10% horse serum, 1 mM sodium pyruvate and antibiotics. RAW267.4 gamma NO(-) cells (ATCC CRL-2278) were a gift from Amnon Bar-Shir (Weizmann Institute of Science, Israel). Cells were cultured in RPMI-1640 supplemented with 10% FBS and antibiotics. All cells were cultured at 37°C with 5% CO_2_ and occasionally checked for mycoplasma infections by PCR using pan-mycoplasma species-specific oligos^66^.

### Plasmids and cell line generation

Unless mentioned otherwise, plasmids were made using restriction-free (RF) cloning^67^. Peroxisome reporter plasmid, pCCL-EGFP-SKL (Addgene #183727), was prepared by inserting human-codon optimized SKL peptide at the C-terminus of the EGFP ORF (before the stop codon) of a lentiviral vector containing EGFP under the control of a Ubiquitin C promoter (UbC-EGFP), using one-step PCR. pRFP-SKL (Addgene #183726) was prepared by subcloning the RFP ORF from a donor pcDNA3-RFP plasmid to replace the GFP in the pEGFP-SKL plasmid. To generate the pHAGE-Pex6-Tomato plasmid (Addgene #183730), *PEX6* ORF was amplified from a pcDNA5-hsPEX6-FRT-Tev-ProtA plasmid (a gift from Ralf Erdmann), *PEX6* 3’UTR was amplified from human cDNA from HEK293T cells, and the Tomato ORF was amplified from a pUbC-tdTomato plasmid (Addgene #183710). Note that during cloning, Tomato was inserted as a single copy and not as a tandem repeat in order to limit the lentiviral genomic size. To construct the 7xHA-Pex6-Tomato plasmid (Addgene #183731), 7xHA was amplified from a 7xHA donor plasmid (a gift from Zvulun Elazar, Weizmann Institute of Science, Israel) and cloned before the start codon of the pCCL-Pex6-Tomato plasmid (**Figure 3A**).

Human HSF1-GFP lentivirus vector was a gift from Maria Vera Ugalde (McGill University, Canada). To create EF1α-HSF1-GFP (Addgene #227820), the UbC promoter driving the expression was replaced by the stronger EF1α promoter, which was PCR-amplified from pTwist-EF1α/Puro plasmid (Twist Bioscience), and subcloned using restriction enzymes AfeI and NotI. Construction of the CRE-MS2/GFP (Addgene #219536) and CRE-MS2/tdMCP-GFP (Addgene #219537) plasmids was done in several steps of restriction digestion-ligation. First, the MS2x24 cassette was subcloned from pNX24-RFP-MS2SL (Addgene #227819) to pHAGE2-FullEF1a-ZsGreen-W (a gift from Yaqub Hanna, Weizmann Institute of Science, Israel) using NotI and BamHI to produce pHAGE-FullEF1a-MS2. Rabbit beta-Globin (RBG) poly A site was amplified from pCAGGSTurbo-cre@1 (gift from Yaqub Hanna) and inserted into the pHAGE2 backbone using BamHI restriction site. The CRE recombinase was amplified from the same plasmid and inserted to pHAGE2 using NotI restriction site. UbC-HA-NLS-tdMCP-GFP and UbC-GFP were PCR amplified from pHAGE-UBC-HA-NLS-tdMCP-GFP (Addgene #40649) and Lenti-UbiC-GFP^11^, respectively, and inserted into the vector backbone using SpeI restriction site. SV40pA was amplified from Lenti-UbiC-GFP by PCR and inserted into the vector using BsRGI restriction enzyme. The entire construct was then transferred to the 3rd generation lentiviral vector pRRLSIN.cPPT.PGK-GFP.WPRE backbone (a gift from Didier Trono, Swiss Federal Institute of Technology Lausanne, Switzerland; Addgene # 12252). Note that during the cloning process, the CRE/GFP construct lost one MS2 repeat by recombination (thus bearing 23 MS2 repeats) and CRE/tdMCP-GFP lost three repeats, thus bearing 21 MS2 repeats.

Lentivirus particles were produced by transiently transfecting the expression plasmid with packaging plasmids, VSVG, RRE and Rev (Addgene plasmids #12259, 12251, and 12253, respectively) into HEK293T cells using TransIT-Lenti transfection reagent (Mirus Bio) or calcium phosphate, and allowed to grow for 72 hrs. The virus-containing media were harvested and concentrated with the Lenti-X concentrator (Clontech), per the manufacturer’s instructions. Viral particles were resuspended in complete DMEM, aliquoted and stored at -80°C until infection. Alternatively, packaging was performed in HEK293SF-PacLV cells^68^ (herein called PacLV cells; a gift from Bernard Massie, McGill University, Canada) according to their provided protocol with minor changes. PacLV cells contain the three genes required for lentivirus packaging under the control of doxycycline-and cumate-inducible promoters. Briefly, PacLV cells were routinely grown in suspension in polycarbonate shaker flasks with filtered caps in HyCell TransFx-H media (Hyclone) supplemented with 4mM glutamine, 0.1% Kolliphor P188 and 1mg/ml penicillin/streptomycin. Cells were incubated at 37°C, 5% CO2 with agitation at 110rpm. For lentivirus packaging, cells were transferred to SFM4-Transfx-293 medium (Hyclone) supplemented with 4mM glutamine and 1mg/ml penicillin/streptomycin. Cells were grown to a concentration of 1.5x10^6^/ml and transfected with 2.67µg plasmid DNA per 1x10^7^ cells using 2µl of PEI-pro transfection reagent (Polyplus) per 1µg DNA. Lentivirus production was induced by adding 1µg/mL doxycycline and 50µg/mL of cumate at 4-6hrs post-transfection and adding 7mM sodium butyrate at 16-24hrs post-transfection. Media was collected 48 and 72hrs post transfection, filtered using 0.45µm membrane, and concentrated by Lenti-X concentrator as described above.

For stable cell line generation, cells (*e.g.* HEK293T, PEX6mut, PEX5^-/-^, HSF1^-/-^ MEFs, MEFs, Ai9 MEFs, EL4, RAW267.4 and CAD cells) were seeded in 6-or 12-well plates and exposed to the viral particles mentioned above (*e.g.* for EF1α-HSF1-GFP, GFP-SKL, RFP-SKL, PEX6-Tomato, 7xHA-PEX6-Tomato, CRE/GFP or CRE/tdMCP-GFP expression) in serum-free media containing 6µg/ml polybrene (Sigma) for 2hrs with occasional shaking, followed by the addition of complete media. Cells with high expression of the fluorescent reporter were selected by FACS sorting (BD Biosciences Aria III). RFP-SKL expressing PEX6mut cells were found to have low viability after FACS sorting and, hence, were not sorted. However, >90% of the cells were RFP-positive after transduction.

### Determining the PEX6 mutation sequence

To identify the mutation in PEX6mut patient-derived fibroblasts, which was not published in the original paper^26^, we extracted DNA (using the Monarch genomic DNA extraction kit, New-England Biolabs) from the cells and performed PCR to amplify all the exons. PCR amplicons sequencing was performed by Plasmidsaurus using Oxford Nanopore Technology with custom analysis and annotation, or by Sanger sequencing at the Weizmann Institute of Science (Israel).

### PEX6mut Complementation Assay

Between 20,000-30,000 PEX6mut-RFP-SKL cells were cultured in a fibronectin-coated 6cm-wide glass-bottom plate (MatTek Corporation). Thereafter, 10,000-15,000 HEK293T-GFP-SKL cells were co-cultured along with the PEX6mut-RFP-SKL cells in the same plate for 3 or 6 days. At the desired time-point, the cells were washed once with PBS and cultured in Leibovitz (L-15) media supplemented with 10% FBS. For the conditioned media and Transwell experiments (**Figure 1E-F**), ∼10,000 PEX6mut-RFP-SKL expressing acceptor cells were cultured alone in a 12-well glass bottom plate (MatTek Corporation), co-cultured either directly with ∼5000 donor cells, or with 5000 donor cells separated using Transwell inserts (pore size of 0.4µm; Corning), or cultured alone with conditioned medium derived from donor cells. Conditioned media was harvested from donor cells, centrifuged for 10min at 500x*g*, and the supernatant was added to acceptor cells cultured separately. Acceptor cells were monitored for peroxisome formation by live imaging in a temperature-controlled chamber and for mRNA transfer by smFISH. Percentage complementation was defined as the number of cells with a punctate pattern of RFP-SKL out of the total number of observed cells.

### HSF1^-/-^ Complementation Assay

WT, HSF1^-/-^ MEFs, HSF1^-/-^ MEFs expressing EF1α-HSF1-GFP, and co-cultures were seeded on fibronectin-coated coverslips in a 12-well plate. After 48hrs co-culture, plates intended for heat shock were wrapped with parafilm and incubated in a 42°C water bath for 1hr. Samples at 37°C (control conditions) were left in the incubator. Following heat shock, samples were immediately taken for FISH analysis.

### CRE Complementation Assay

Ai9 MEFs were co-cultured at a 1:1 ratio with HEK293T, WT MEFs, EL4, RAW267.4 or CAD cells expressing GFP, CRE/GFP or CRE/tdMCP-GFP for 72hrs on fibronectin-coated glass bottom 12-well plates (MatTek), 8-chamber slides (ibidi) or 96-well Round Glass Bottom µ-Plates: #1.5H (ibidi). Transwell experiments in 12-well glass-bottom plates were performed as described above for PEX6mut cells. Live imaging to score the number of tdTomato-positive (red) cells was done using the 10X objective on a Zeiss AxioObserver Z1 imaging system equipped with an Illuminator HXP 120-V light source and a Hamamatsu Flash 4 sCMOS camera. Prior to live imaging, the media was replaced with Leibovitz media (L-15) supplemented with 10% FBS. In some experiments, 0.5µg/ml Hoechst 33342 was added as well. The number of red cells in each well were counted and the percentage calculated based on the number of initial Ai9 MEFs seeded at the start of the experiment.

### smFISH/smFISH-IF

A library of 48 Cy5-labeled FISH probes (20-mers) to detect Tomato mRNA was a gift from Shalev Itzkovitch (Weizmann Institute of Science, Israel) and were previously described^10^. Quasar 670-labeled probes for mouse HSPA1A (HSP70)^69^ were a gift from Robert H. Singer (Albert Einstein College of Medicine, NY, USA). Quasar570-labeled probes for human HSF1 are listed in **Supplementary File 1**. Cy5-labeled probes for MBS were previously described^11^. smFISH was carried out according to a previously described protocol^70^ without any modifications. For simultaneous analysis of mRNA and HA-tagged proteins, coverslips were fixed, permeabilized, pre-hybridized and hybridized with FISH probes as per the smFISH protocol. After the hybridization step, cells were washed twice with pre-hybridization buffer for 15min each at 37°C, rinsed three times in 2xSSC. Thereafter, the coverslips were blocked with blocking buffer (2% BSA in PBS supplemented with 1% Tritox-X-100 and 10U/mL of RNAsin) for 1 hour with shaking. After blocking, the cells were treated with a mouse anti-HA primary antibody (Biolegend, Clone: 16B12; Dilution: 1:1000) for overnight at 4°C in blocking buffer. After three washes with PBS for 5min each, coverslips were treated with AlexaFluor488-tagged rabbit anti-mouse secondary antibody (Jackson Immunoresearch; Dilution: 1:800) in blocking buffer for 1hr at room temperature in the dark. After three quick washes with PBS for 5min each, samples were stained with DAPI (0.5 µg/mL in 2x SSC) and washed with 2xSSC for 5min. The coverslips were then mounted on an imaging slide using Prolong Glass antifade reagent (Molecular Probes). Slides were sealed with nail polish (Electron Microscopy Sciences), dried at room temperature for 24-48hrs, and stored at -20°C until imaging. smFISH/FISH-IF images were captured using a Zeiss AxioObserver Z1 DuoLink dual camera imaging system equipped with an Illuminator HXP 120-V light source, Plan Apochromat 63x or 100x 1.4 NA oil-immersion objective, and a Hamamatsu Flash 4 sCMOS camera. Thirty 0.2μm step *z*-stack images were taken for each field using a motorized *XYZ* scanning stage 130 × 100 Piezo, and ZEN2 software at either 0.0645 µm per pixel (100x) or 0.130 µm per pixel (63x).

The number of cytoplasmic mRNA FISH spots, transcription sites, and nascent mRNAs per transcription site were scored using FISH-Quant^71,72^. For image presentation, representative cells from each condition were minimally adjusted for brightness and contrast using Fiji^73^.

### Immunofluorescence (IF)

To check for the expression and localization of PEX14 or Catalase proteins by IF, cells grown on coverslips were fixed and permeabilized as described for FISH. Samples were then blocked (3% BSA, 0.1% Tween in PBS; PBST) for 1hr. Samples were then incubated with primary antibodies rabbit anti-PEX14 (Proteintech Cat #10594-1-AP; a gift from Orly Laufman, Weizmann Institute of Science, Israel; Dilution: 1:200) or Rabbit anti-catalase (gift from Orly Laufman; Dilution: 1:200) in PBST+BSA for 2hrs at room temperature, washed 3 times in PBST, then incubated with the secondary antibody AlexaFluor647-goat anti-Rabbit IgG (Jackson Immunoresearch; Dilution: 1:400) in PBST+BSA for 45 min. Samples were washed as before, incubated with 0.5µg/ml DAPI in PBS for 2min, washed once in PBS, mounted on slides using Prolong Glass, and dried for 24 hrs. IF samples were imaged using the Zeiss AxioObserver Z1 microscope using the 63x objective and a single *z* plane as described above. Images were processed and adjusted using FIJI.

### TNT staining and quantification

For TNT staining, cells were grown on coverslips in co-culture for 24hrs, then fixed in 3% paraformaldehyde + 1% glutaraldehyde in PBS for 10 min, followed by quenching with 0.1M glycine in PBS for 10min. Samples were washed twice in PBS, permeabilized in 0.1% Triton x-100 in PBS for 10 min and washed twice with PBS. Samples were incubated with 50nM phalloidin-TRITC (Sigma) for 30min at RT. Samples were washed twice with PBS, incubated with 0.5µg/ml DAPI in PBS for 2min, washed once in PBS, then mounted on slides using Prolong Glass. TNT samples were imaged using the Zeiss AxioObserver Z1 microscope using the 63x objective as described above. Nineteen 0.5μm step *z*-stack images were taken for each field. Images were processed and adjusted using FIJI. TNTs were counted manually. Only TNTs connecting donor to acceptor cells were counted (as opposed to donor-donor and acceptor-acceptor TNTs). TNTs were defined as thin connections (typically <0.5µm in width) that connect cells in a straight line. Donors were identified based on GFP expression and/or cell or nuclear morphology. Between 100 to 200 donor cells were counted in 8-11 fields of view per sample.

### Puromycinylation -Proximity Ligation Assay

To detect the *in situ* translation of transferred HA-PEX6-Tomato mRNAs, 50,000 donor (HEK293T stably expressing 7xHA-Pex6-Tomato) and 100,000 acceptor cells (PEX6mut) were cultured either alone or together on fibronectin-coated coverslips in 12-well plates. After about 2 days of co-culture (∼70-80% confluence), cells were treated with 2µg/mL of Puromycin (Sigma) for 15min at 37°C. The cells were then rinsed three times in PBS supplemented with 5mM MgCl_2_ (PBSM), with 4% Paraformaldehyde (Electron Microscopy Sciences) and quenched with 0.1% Glycine (Sigma) in PBSM. After 2 washes of 10min each with PBSM, samples were permeabilized with 0.1% Triton-X-100 (Sigma) in PBS for 10min. The coverslips were blocked with blocking buffer (2% BSA in PBS supplemented with 1% Triton-X-100 and 10U/mL RNAsin) for 1hr with shaking. For primary antibody incubation, cover slips were incubated with mouse anti-Puromycin antibodies (Sigma Cat # MABE343; Dilution: 1:10,000) and rabbit anti-HA antibodies (Cell Signaling Clone C29F4; Dilution: 1:1000) in a humidified hybridization chamber as mentioned in the smFISH protocol. The reaction volume was 40µl and the incubation was done at 4°C. After primary antibody incubation and three quick washes with PBS, the Proximity Ligation Assay (PLA) was carried out as per the manufacturer’s instructions (Sigma) using the rabbit PLA Plus and mouse PLA Minus antibody and a far-red detection kit. After the last recommended wash with Buffer B, coverslips were stained with DAPI (0.5 µg/mL in PBS), then washed with PBS for 5min. The coverslips were then mounted on a slide using Prolong Glass antifade reagent (Molecular Probes). Slides were sealed with nail polish (Electron Microscopy Sciences) and stored at -20°C until imaging. Images were captured with Zeiss AxioObserver Z1 microscope as described for smFISH above.

### Quantitative Reverse Transcription-Polymerase Chain Reaction

Total RNA was extracted from cells using Nucleospin RNA Mini kit (Macharey Nagel). Between 2-5µg of RNA was depleted of DNA using PerfecTa DNAse I kit (Quantbio, Beverly, MA) according to the manufacturer’s instructions. The time of DNAse I incubation was extended from 30min to 2hrs. DNA-depleted RNA was then reverse transcribed using the qScript first strand cDNA synthesis kit (QuantaBio, Beverly, MA). qPCR for *PEX6* cDNA was performed using LightCyler 480 device (Roche Diagnostics, Basel) and Power SYBR green mastermix (Applied Biosystems) at Tm of 60°C.

### Western blot

To detect CRE recombinase protein levels, HEK293T cells expressing CRE/GFP or CRE/tdMCPGFP were harvested and protein was extracted using RIPA buffer. Then, 1mg of protein per sample was separated by SDS-PAGE (4-10%) electrophoresis and CRE protein was detected using rabbit anti-Cre-Recombinase antibodies (Synaptic Systems; Cat# 257 003). For loading control, mouse anti-Vinculin (Sigma cat# V9131) was used.

### Statistical Analysis

Comparisons involving two experimental conditions were done using a one-tailed *t*-test and while comparisons involving >2 conditions were done using ANOVA, followed by pairwise multiple comparisons using the Tukey test. Adjusted P<0.05 was considered statistically significant. All indicated calculations were performed by GraphPad Prism software (GraphPad Software, Inc.).

## Acknowledgments

We thank Ruth Scherz-Shouval (Weizmann Institute of Science, Israel), Shalev Itskovitz (Weizmann Institute of Science, Israel), Ralf Erdmann (Ruhr Universitat Bochum, Germany), Maria Vera Ugalde (McGill University, Canada), Robert H. Singer (Albert Einstein College of Medicine, NY, USA) and Steffen Jung (Weizmann Institute of Science, Israel) for cells and reagents. Miriam Waghalter created the EF1α-HSF1-GFP plasmid. We thank the FACS unit staff at the Weizmann Institute of Science for their help. This work was funded by grants to J.E.G. from the Weizmann Institute of Science, including from the BINA center, the Joel and Mady Dukler Fund for Cancer Research, the Jean-Jacques Brunschwig Fund for the Molecular Genetics of Cancer, a Proof-of-Principle Grant from the Moross Integrated Cancer Center, and the Kekst Family Institute for Medical Genetics. J.E.G. holds the Besen-Brender Chair of Microbiology and Parasitology.

## Author contributions

G.H., S.D., and A.G-R. performed the experiments; G.H. and J.E.G. designed the experiments and supervised the project. S.D., G.H., and J.E.G. wrote and edited the manuscript. J.E.G. secured funding.

---

**Supplementary Figure 1.**
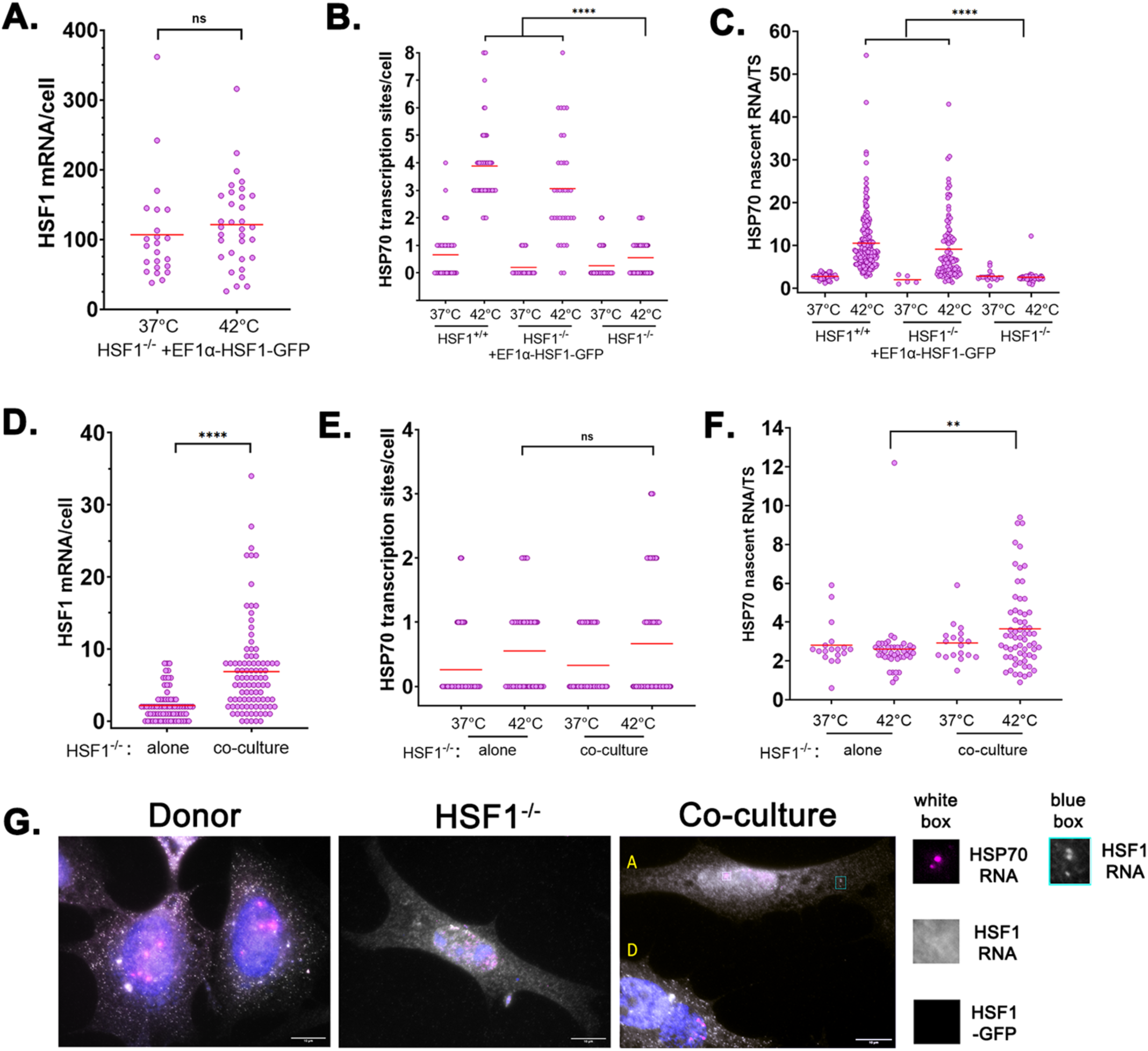
Complementation of the heat shock response in HSF1^-/-^ cells by intercellular mRNA transfer. **A**. FISH analysis of HSF1-GFP mRNA expression (mRNAs/cell) in donor cells. In this and all other panels, each spot on the graph represents a single cell; red line is the average. **B.** FISH analysis for the number of HSP70 transcription sites (TSs) per cell in wild-type (HSF1^+/+^) MEFs, HSF1^-/-^ MEFs or donor HSF1^-/-^ MEFs over expressing HSF1-GFP. Note that cells are tetraploid and during DNA duplication can yield up to 8 TSs. **C.** FISH analysis of the calculated number of HSP70 nascent RNAs per TS in wild-type (HSF1^+/+^) MEFs, HSF1^-/-^ MEFs or donor HSF1^-/-^ MEFs over-expressing HSF1-GFP. Note that the small number of HSF1^-/-^ cells at 37°C is because only a few cells had detectable TSs. **D.** FISH analysis of transferred HSF1-GFP mRNA in acceptor cells cultured alone or in co-culture with HSF1^+/+^ donors. **E.** FISH analysis for the number of HSP70 transcription sites per cell in acceptor cells cultured alone or in co-culture with HSF1^+/+^ donors. **F.** FISH analysis of the calculated number of nascent HSP70 RNAs per TS in acceptor cells cultured alone or in co-culture with HSF1^+/+^ donors. **G.** FISH images of donor with HSF1^+/+^ donor cells (left), an HSF1^-/-^ acceptor cell alone (middle) and HSF1^-/-^ acceptor cell (yellow ‘A’) in co-culture with a donor cell (yellow ‘D’) (right) after 1 hr of heat-shock. Blue – HSF1-GFP protein. Gray – FISH for HSF1-GFP mRNA. Magenta – FISH for HSP70 mRNA. White box – enlarged images of the boxed TS in all three channels are shown on the right. Cyan box – enlarged image of transferred HSF1-GFP mRNA spots in the acceptor cell is shown on the right.

**Supplementary Figure 2.**
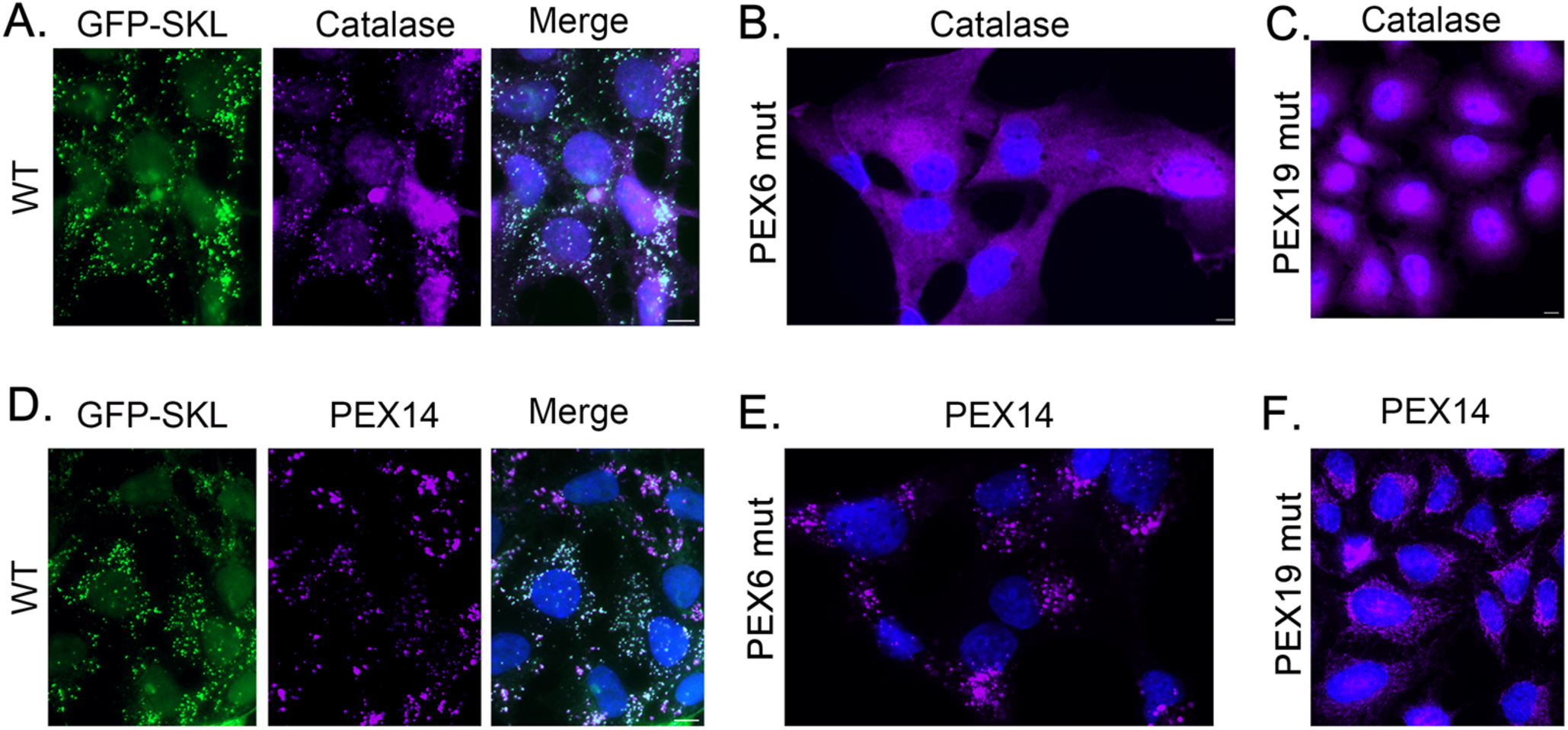
Functional peroxisomes and peroxisome ghosts. **A.** HEK293T (WT) cells expressing GFP-SKL (green), stained for catalase by IF (magenta), and labeled with DAPI (blue). *Merge –* merged panels. Scale bar – 10µm. Note presence of co-localized puncta (*i.e.* functional peroxisomes). **B.** and **C.** PEX6mut and PEX19mut (PEX19^-/-^) cells, respectively, stained for catalase by IF (magenta) and labeled by DAPI (blue). Note absence of catalase-labeled puncta. **D.** HEK293T (WT) cells expressing GFP-SKL, stained for PEX14 by IF (magenta), and labeled by DAPI. Note presence of co-localized puncta (*i.e.* functional peroxisomes). **E.** and **F.** PEX6mut and PEX19mut cells, respectively, stained for PEX14 by IF (magenta) and labeled by DAPI. Note presence of PEX14, but not catalase, puncta (as shown in panels B. and C., *i.e.* non-functional peroxisome ghosts).

**Supplementary Figure 3.**
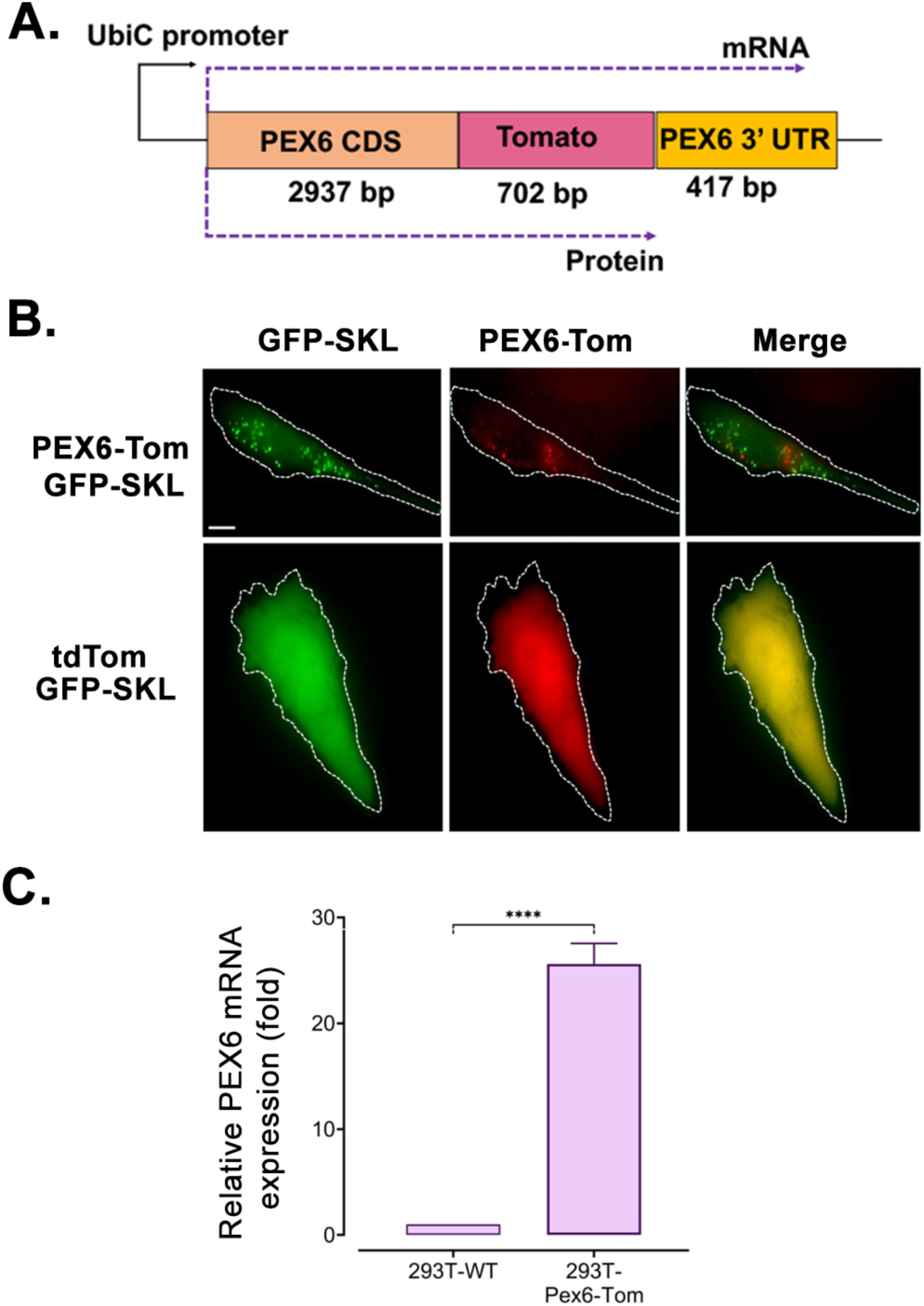
Generation of a PEX6 overexpression construct and cell line. **A.** Schematic of PEX6-Tomato. The Tomato ORF was inserted between the CDS and 3’UTR of human PEX6 and in-frame to PEX6 to produce a PEX6-Tomato fusion protein, as indicated. The construct was expressed under a constitutive ubiquitin promoter. Lengths of each segment are marked in base pairs. **B.** PEX6-Tomato can rescue a peroxisome mutant phenotype. PEX6mut cells were transiently co-transfected with plasmids expressing either PEX6-Tomato and a GFP-tagged peroxisomal marker, GFP-SKL, or a control construct expressing tdTomato alone along with GFP-SKL. Images were taken 72hrs post transfection. Scale bar: 10 µm. **C.** Over-expression level of PEX6-Tomato mRNA. Stable PEX6-Tomato HEK293T cells were created by lentiviral infection. PEX6 and PEX6-Tomato gene expression was quantified by RT-qPCR. 18S gene expression served as endogenous control. Data is shown as the mean of three replicates and the corresponding SEMs. ****: P<0.0001

**Supplementary Figure 4.**
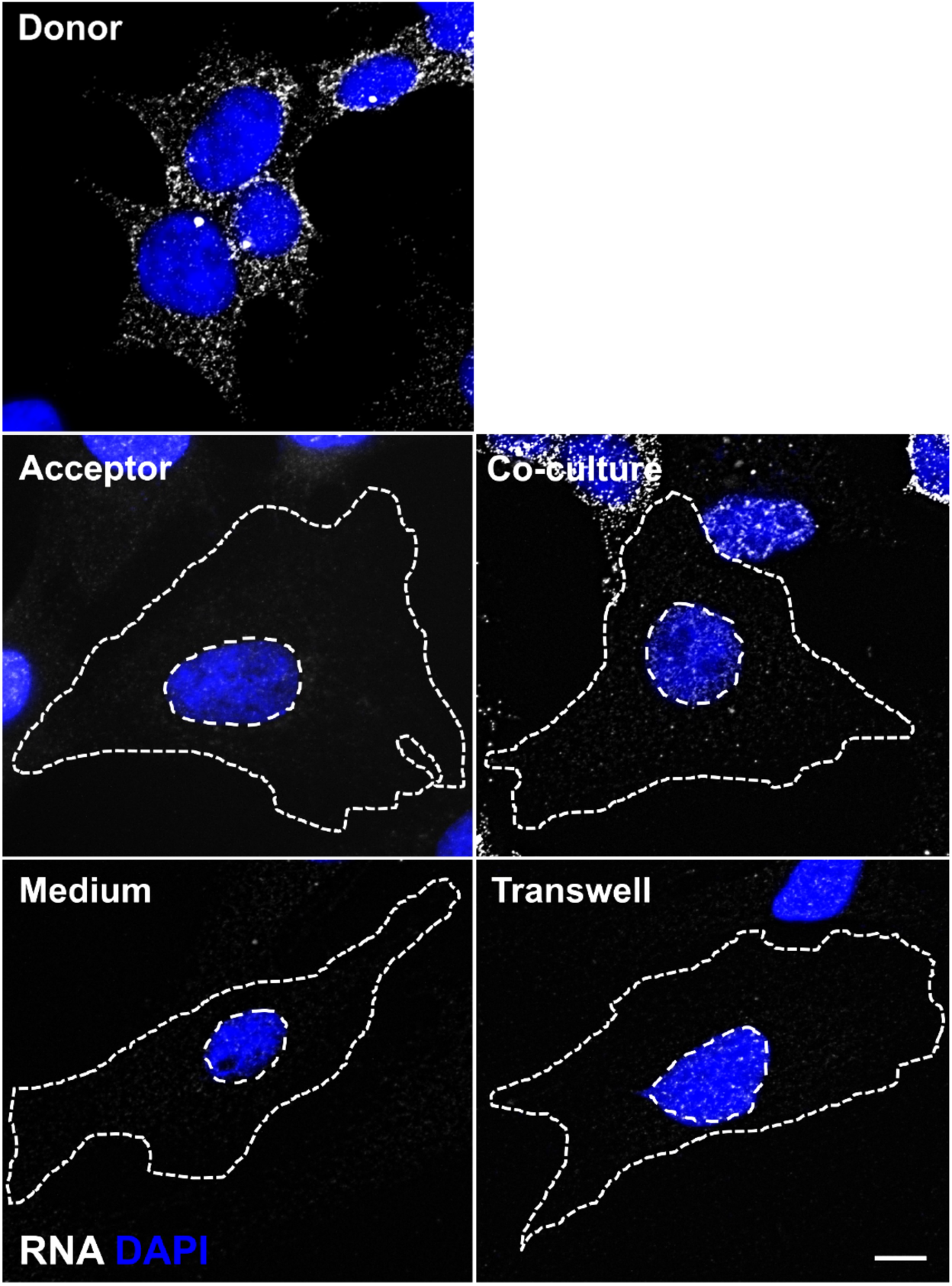
Representative smFISH images. Representative smFISH images of PEX6-Tomato mRNA (white spots) in donor HEK293T cells, PEX6mut acceptor cells cultured alone (Acceptor), PEX6mut acceptor cells in co-culture with donor cells, and PEX6mut acceptor cells cultured either with donor cell-derived conditioned media or co-cultured in a Transwell setup. Cells were labeled with DAPI (blue). Dashed lines outline the approximate cell and nuclear boundaries. Scale bar - 10 µm. Image of the donor cells was deconvoluted using FISH-Quant.

**Supplementary Figure 5.**
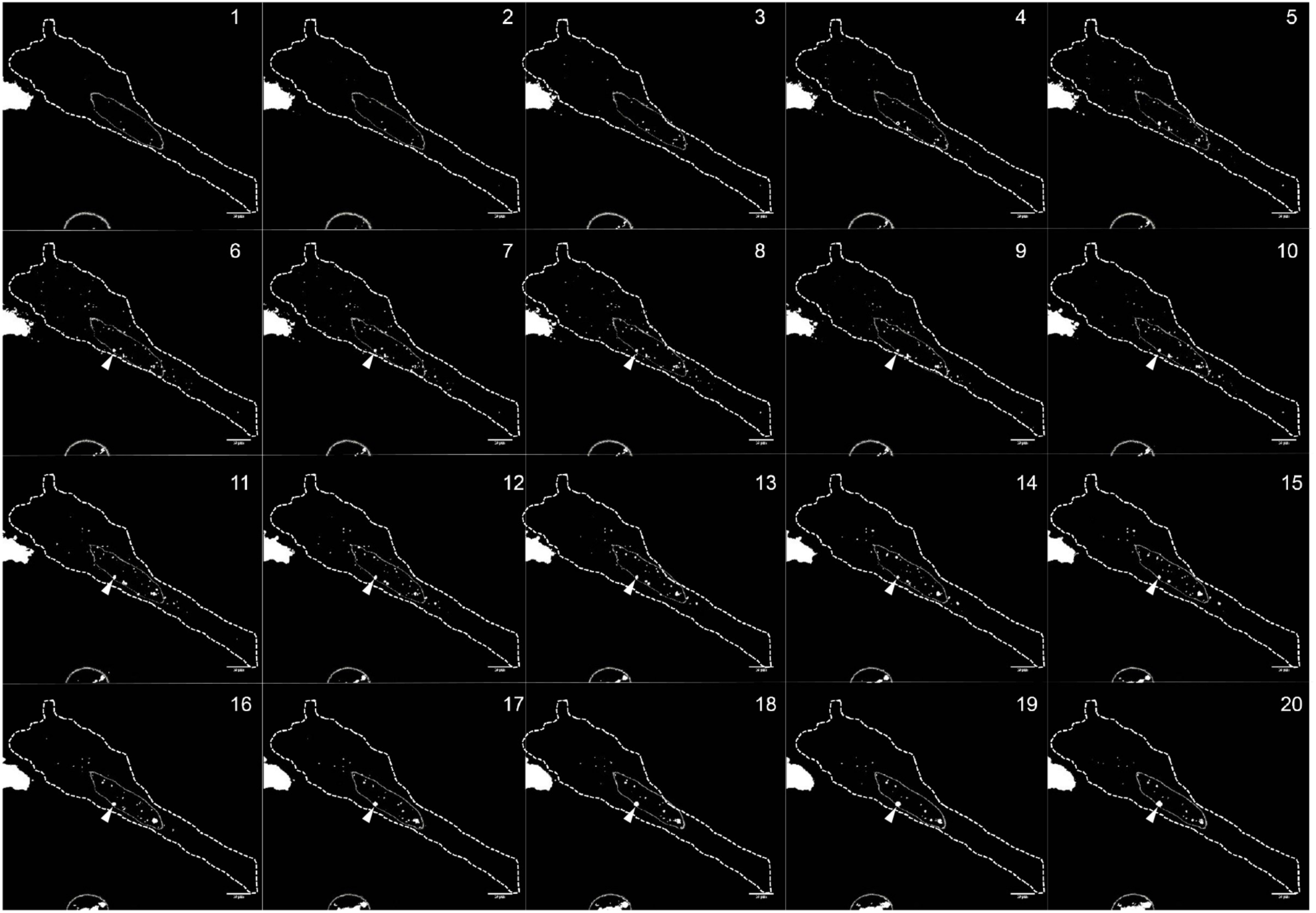
Distribution of Puro-PLA spots across a PEX6mut acceptor cell in co-culture. Twenty 0.2-μm step *z*-section images of a representative PEX6mut acceptor cell with the nascent polypeptides of HA-Pex6-Tomato as detected by Puro-PLA is shown, as described in Figure 3I. The images were thresholded using ImageJ to detect only the PLA spots and the outline of nuclei for each section. The number on the top left corner corresponds to the section number from the basal end to the apical end. Based on the intensity of nuclear staining by DAPI, Sections 5-11 encapsulate most nuclear volume. Dashed outlines indicate the approximate cell borders. As shown, nuclear/perinuclear PLA spots are mainly found in the higher *z* sections (*e.g*. 11-20), as compared to the lower ones (*e.g*. 1-10). A representative non-specific spot that is persistent through multiple *z*-sections is marked with an arrowhead. Scale bar -10 µm.

**Supplementary Figure 6.**
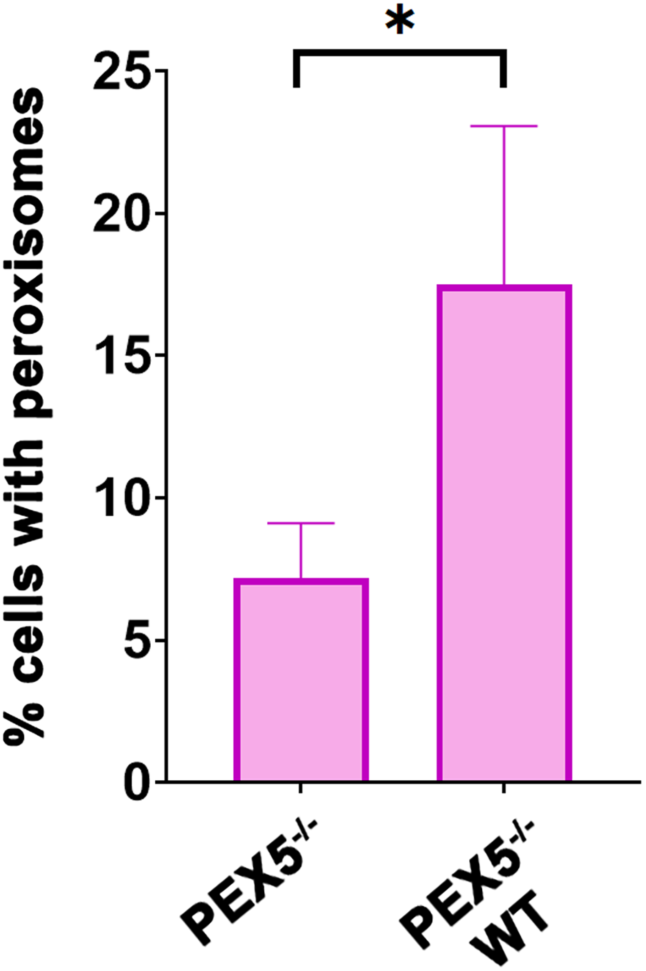
Complementation of PEX5^-/-^ cells in co-culture with WT cells. A bar graph showing the percentage of PEX5^-/-^ RFP-SKL cells bearing red puncta in single cultures or in co-cultures with WT cells after six days. An average of four biological repeats is shown. Error bars indicate standard deviation. * - p<0.05.

**Supplementary Figure 7.**
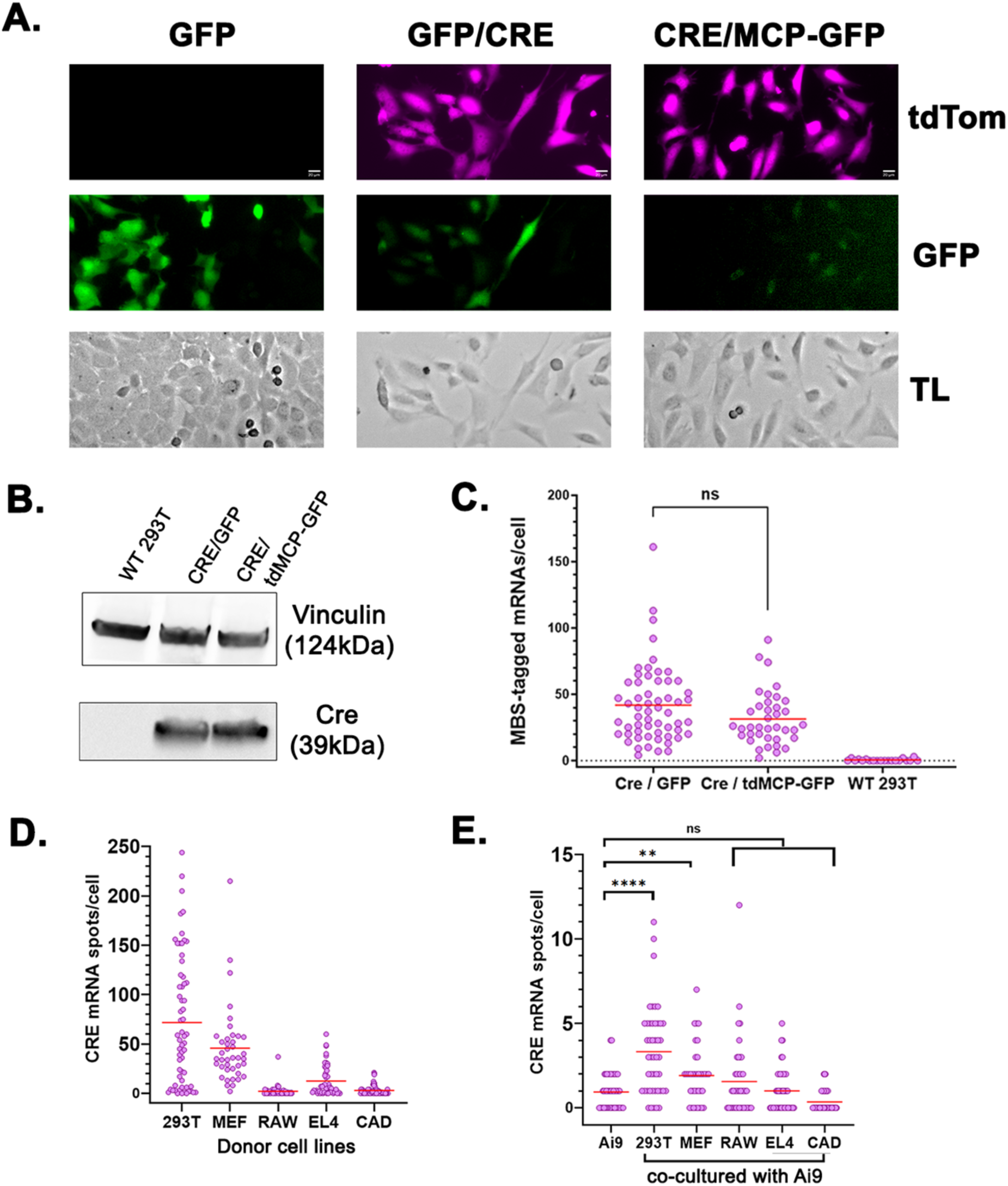
CRE expression in donor cells. **A.** Transient transfection of Ai9 MEFs with CRE/GFP or CRE/MCP-GFP, but not GFP, results in tdTomato expression. Shown are images of Ai9 cells in the red channel (tdTomato), green channel (GFP) and transmitted light (TL) four days post transfection. Note that the GFP expression of CRE/GFP and CRE/tdMCP-GFP is low compared to GFP alone. **B.** CRE protein from CRE/GFP and CRE/MCP-GFP is expressed at similar levels in transduced HEK293T cells, as detected by Western blot. **C.** CRE mRNA from CRE/GFP and CRE/MCP-GFP is expressed at similar levels in transduced HEK293T cells, as detected by smFISH. Each dot in the graph is a single cell. **D.** Comparison of CRE mRNA expression in the different donor cells, as detected by smFISH. Each dot in the graph is a single cell. **E.** Comparison of CRE mRNA transfer to Ai9 cells from the different donor cells after 24hr of co-culture, as detected by smFISH. Each dot in the graph is a single cell. ** - p<0.01 **** - p<0.0001

**Supplementary Figure 8.**
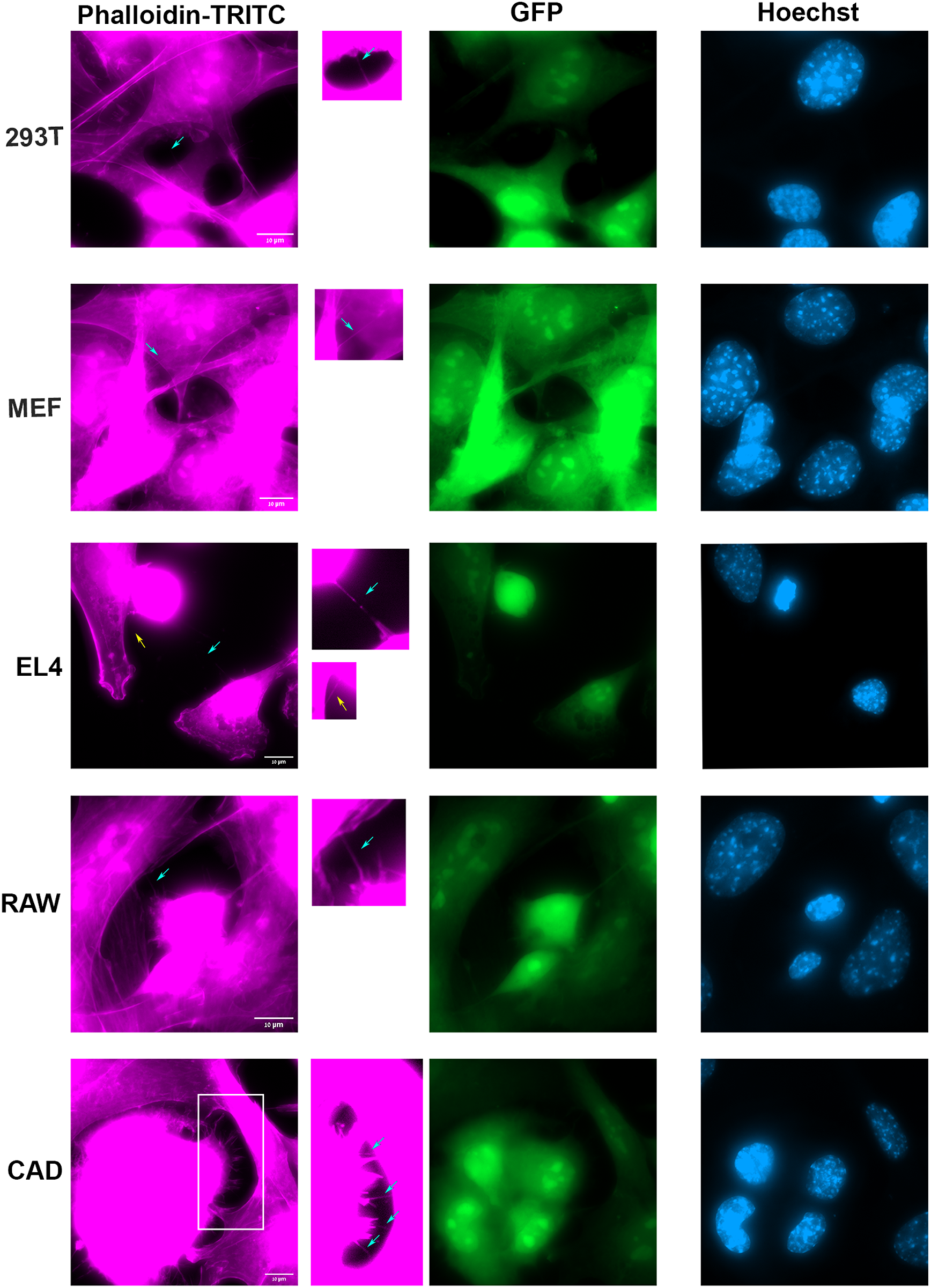
Donor and acceptor cells are connected by TNTs. Images of cytoskeleton staining (Phalloidin-TRITC, magenta), GFP expression (green) and Hoechst (blue) for TNT-connected cells are shown. Note that donor cells are GFP-positive. TNTs are difficult to detect and require increasing the signal intensity of selected regions in the image. TNTs are indicated by arrows. Maximum projections of the *z-*stack images are shown.

**Supplementary File 1. List of HSF1 FISH probes**

